# From plant populations to communities: using hierarchical trait environment relationships to reveal within ecosystem filtering

**DOI:** 10.1101/825471

**Authors:** Lucas Deschamps, Raphaël Proulx, Nicolas Gross, Guillaume Rheault, Vincent Maire

## Abstract

Explaining the existence of highly diverse plant communities under strong abiotic filtering is a long-standing challenge in ecology. Hierarchical aspects of abiotic and biotic filters are rarely taken into account and studies focus mainly on community-level aggregated patterns. Because variations in biotic conditions might take place in short abiotic gradient and within the tolerance of species in regional pool, it is likely that biotic filtering will select individuals within species and adjust population characteristics. To challenge this idea, we replicated a diversity gradient in four highly contrasted wetlands with an almost complete species turn-over, sampling individuals in communities irrespective of their taxonomic identities or status. Using hierarchical distributional modelling, we analyzed the variation of the mean and dispersion of functional trait space at the ecosystem, community and species level. We found that the abiotic differences between ecosystems filtered species contrasted in their growth/nutrient conservation trade-off, while within ecosystems community variation were mainly due to the partitioning of canopy and leaf adaptations to light conditions. We found strong species-specific functional and demographic responses of dominant species along the diversity gradient, especially for traits linked to biomass and space occupation. Two contrasted strategies emerged, with species using plasticity to maintain equally dense populations, while others used plasticity to become overwhelmingly abundant when in favorable conditions. Our results demonstrate that within ecosystems, variation in biotic conditions selects individuals within populations, revealing the importance of phenotypic variation for a species to be maintained in more or less diverse communities. Because phenotypic variations are related to demographic responses, it offers a way to link the study of species diversity and eco-evolutionary dynamics.

## Introduction

Environmental selection of individuals growing in plant communities is described as a hierarchical succession of abiotic and biotic filters (Lortie *et al.*, 2004; HilleRisLambers *et al.*, 2012). Consequences of these filters have been greatly unraveled using functional traits, which are measurable characteristics of individuals linked to their fitness (Violle *et al.*, 2007). When aggregated at the community level, the response of trait values to environmental gradients is the most widely used tool to reveal how different environments filter individuals for their ability to exhibit traits allowing them to grow under a given set of conditions and resources (thereafter summarized as “ecological constraints”). The *directionality* of the filtering describes the displacement of the mean value of the trait to cope with the set of ecological constraints occurring along an environmental gradient. The filtering *intensity* occurs when constraints at a given position along the environmental gradient reduce or enlarge the envelope of trait values allowing individuals to grow and reproduce (Laughlin & Joshi, 2015). The former process is tackled with the community-aggregated average, while the latter is studied with the statistical dispersion (thereafter called “dispersion”). Variation in community mean and dispersion might be due to species sorting, change in species relative abundances or even intra-specific responses (De Bello *et al.*, 2012; Bjorkman *et al.*, 2018). This trait-environment approach successfully allowed understanding the main abiotic drivers of functional traits (Maire *et al.*, 2015), predicting species turn-over across ecosystems (Shipley, 2010; Maire *et al.*, 2012), and revealing the variation of principal biotic mechanisms in play in different ecosystems characterized by contrasted abiotic conditions (Hulshof *et al.*, 2013; Berdugo *et al.*, 2018). Because filtering acts on individuals, it remains unclear if the information carried by community level traits is sufficient to understand the constraints driving trait assembly, especially within ecosystems where abiotic gradients might not be sufficient to exceed species tolerance.

Within ecosystem, local biotic constraints (e.g., intra- and inter-specific competition, facilitation, enemy-release) also shape the direction and intensity of filtering. At this scale, the relative importance of abiotic filtering likely decreases in comparison with biotic filtering (Chalmandrier *et al.*, 2017). Directionality may occur when relative fitness advantage of species vary, displacing the community trait mean toward the most competitive strategy in case of competitive hierarchy (Kunstler *et al.*, 2012), or toward the center of a multimodal distribution when different equivalent strategies coexist (Le Bagousse-Pinguet *et al.*, 2016). Filtering intensity will decrease when the dispersion of trait values is increased by competitive exclusion (Mayfield & Levine, 2010), or by limit-to-similarity process forcing plants to exploit different strategies to balance intra- and inter-specific competition (Grime, 2006; Gross *et al.*, 2013). To study the response of directionality and intensity to the strength of biotic filtering, one would require an environmental gradient that is methodologically feasible, biologically sound and independent from other environmental gradients. Rarely these considerations have however been applied to study environmental selection in natural ecosystems.

For a given community trait distribution, filtering may still operate at the species level. Both directionality and intensity of filtering may be used to describe the niche of species (Austin & Smith, 1989). Individuals of different populations may tend to exhibit different mean trait values depending upon the competitive advantage it provides (Weiner & Thomas, 1986). The dispersion of species traits may be contracted, when individuals tend to exploit the same specialized resources and limit overlap with neighbors (Violle *et al.*, 2012; Hulshof *et al.*, 2013; Meilhac *et al.*, 2019), or dilated when neighbors vary in strategies, potentially avoiding competition between very similar conspecific or to limit between species differences in fitness (Le Bagousse-Pinguet *et al.*, 2014).

Such changes in species trait distribution may have two demographic consequences, as described by Richards *et al.* 2006. *Jack-of-all-trade* strategists described species that exploit their intra-specific variations to maintain their demography through contrasted environmental constraints. Such species will be as dense when dominating a community as when coexisting in diverse ones. *Master-of-some* strategists would instead take advantage of more favorable environment to dominate and maintain higher demographic rates, and will be denser in dominated communities than in more diverse ones. While it has already been demonstrated that accounting for intraspecific trait variation allowed to detect community level patterns potentially linked to biotic interactions (Siefert *et al.*, 2015; Chalmandrier *et al.*, 2017), few studies explored the way intra-specific variation might shape community functional structure.

Directionality and intensity needs to be studied on multiple traits that describe ability of individual to cope with changes to environmental (biotic and abiotic) constraints. It has been shown that variation of distinct traits takes place at different levels of biological organization (de Bello *et al.*, 2013; Siefert *et al.*, 2014), and that uncorrelated sets of functional traits rely to different resources and filtering processes (Maire et al. 2012, Maire et al. 2009, Pontes et al. 2015). Indeed, the multidimensionality of their ecological niches allows individuals to be adapted to multiple environmental constraints (Cornwell *et al.*, 2006). To better understand species selection within communities, we need therefore to select a set of traits representing the different functional niche dimensions of individuals.

To explore consequences of successive abiotic and biotic filters on species assemblages, we designed an original hierarchical approach that disentangles consequences of filtering at the species, community and ecosystem levels. We first selected wetland ecosystems characterized by highly contrasted site pH, but within the same climatic envelope. Within each ecosystem, we selected natural communities along a species diversity gradient, from mono-dominated to highly diverse communities, which minimized for differences in abiotic conditions (Rheault *et al.*, 2015). Because diversity gradient is known as the result of different modalities of species interactions (Chesson, 2000; Levine & HilleRisLambers, 2009) and resource partitioning (Tilman *et al.*, 1997b), and is consequently thought as an important driver of ecosystem functioning (Tilman *et al.*, 1997a), we choose it to describe potential variations in biotic filtering. Plant ramets were sampled irrespective of their taxonomic identity to both study the community distribution and capture the role of phenotypic variation in the local selection processes. We formulated models with explicit parameters for mean and dispersion of community and species distribution of key functional traits known to be related to nutrient immobilization and light acquisition and processing. Accounting for differences between ecosystems, we tested through modeling of community trait distribution (1) where biotic filtering could be more accurately detected: community or species level, or said differently, what is the role of intra-specific variation in fine-scale community assembly; (2) if the response is trait-dependent; and (3) if species exhibited contrasted growth strategies in response to biotic filtering, linking functional trait responses to population fitness variation. Along the pH gradient, we assumed that traits linked with nutritive stress response (e.g. LDMC, SLA) would follow a directional filtering at community level, displacing community mean and dispersion. In contrast, along the diversity gradient at species level, we expect traits linked with competitive ability for light capture and space occupation strategy (e.g. LA, EL) to respond stronger than other traits.

## Material and Methods

### Study sites

In the lowland of the Fleuve St-Laurent in Eastern Canada, a diversity gradient of eight communities were replicated in four highly contrasted wetland ecosystems (total number of communities = 31, with only seven plots sampled in the wet meadow). Every ecosystem was situated in the same climatic envelope (mean annual temperature = 5.4°C, mean annual precipitation = 1030.5mm, mean length growing season = 112 days) and within an acidic regional context due to the proximity to the granitic Canadian shield. These ecosystems are ranked along a fertility gradient (soil pH as proxy, Fig. S1a), and characterized as bog (Lac-à-la-Tortue, 46°33’15"N 72°39’46"W), fen (Red-Mill, 46°25’38.9"N 72°29’46.6"W), wet meadow (SCIRBI, conservation society, 46°04’12.9"N 73°10’11.1"W) and fluvial marsh (Maskinonge, 46°11’39.1"N 72°59’58.7"W). Within each site, we selected communities of similar area to build a species richness gradient. The gradient ranged from two species to highly diverse community (up to 16 species, Fig. S1b) and were of comparable range between sites (Fig. S1b). Importantly, there was no relationship between soil pH nor ramets density and species diversity of communities within and among ecosystems (Fig. S1c and d), and the design has been thought to minimize abiotic differences within ecosystems (Rheault et al. 2015).

### Vegetation sampling

Within each community, we sampled 80 individuals, irrespective of their taxonomic characteristics, with at least two mature leaves and during two sampling campaigns (from the 14^th^ of June to the 5th of July and from the 22th of August to the 3^rd^ of September 2016). The method of point-plant distance sampling was used and simultaneously realized by two independent harvesters, each one directed by successive random bearings and distances. When approaching from the edge of the community, the harvester bounced with an angle of 45º toward the plot. At the point determined by those indications, the closest mature plant was harvested, and its distance to the point measured. Plant density within communities were computed using the following equation, where *dens*_*p*_ is the density in individuals/m^2^ of plot *p*, *n* the number of individuals harvested in the plot, and *d*_*pi*_ the distance, in cm, between the point and the individual *i* of plot *p*:

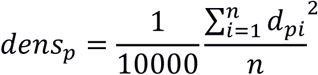

### Trait measurements

We selected a set of plant functional traits that are linked to different biological functions and covary along different dimensions of the species niche (Fig. S2). Leaf Area (LA) and Extended Length (EL) and leaf angle are linked to the space occupation strategies that are used to compete for light interception (Hikosaka & Hirose 1997, Weiner & Thomas 1986), while superficial chlorophyll content represents the fine scale adaptation to optimize light utilization by the leaf (Kull & Niinemets 1998). All these traits are also components of the leaf energy balance that regulates leaf temperature and biochemical processes. Leaf dry matter content (LDMC) and flavonoid content are related to nutrient conservation, involved in response to stress and/or herbivory (Hodgson *et al.* 2011, Izaguirre *et al.* 2007). Specific Leaf Area (SLA) directly scales with the relative growth rate of individuals in herbaceous ecosystems (Garnier *et al.*, 2004; Poorter *et al.*, 2012). LA, LDMC and SLA were measured following the protocols described in (Pérez-Harguindeguy *et al.*, 2013), on the last mature leaf of plants. EL is the length of individuals from the ground to the edge of their deployed leaves, in cm. Superficial chlorophyll content is the concentration of chlorophyll in the leaf epidermis (μg cm^−2^), and flavonoid content is an index of flavonoids concentration in this superficial layer, which is related to phenol accumulation and UV protection. Both were measured using a portable Dualex device (Force-A, Orsay, France), which uses a combination of fluorescence signals at various excitation band to quantify pigments and chemical compounds. This method have been successfully used to follow the phenology of leaf and individuals (Mattila *et al.*, 2018), as the response of leaf metabolism to nutrients (Scogings, 2018) or light manipulation (Agati *et al.*, 2011). The relationships between superficial chlorophyll content and total chlorophyll extracted with methanol are presented by Fig. S3.

### Data analysis

Our aim is to explain how plant trait varies along environmental gradients and test if the variation of the mean and dispersion of trait values observed at community and species levels is related to differences in the number of species present in the community. We explored this principle across a series of traits to evaluate if the filtering has differently acted on traits.

The bayesian distributional modeling framework has been used to model the community and species trait distribution (Rigby & Stasinopoulos, 2005). This framework allows modeling each parameter of a given probability trait distribution by an independent equation, thus relaxing the fixed dispersion assumption classically stated in the GLM framework (Smyth, 1989; Cepeda-Cuervo, 2015). We modeled the distribution of traits at the community and the species level with two parameters distributions. *Gamma* distribution parameterized in terms of mean (*μ*) and dispersion (*ϕ*) have been used to model traits with strictly positive distribution (all but LDMC). As being distributed on [0,1] interval, LDMC has been modeled as a *beta* distribution parameterized by the mean (*μ*) and precision (*ϕ*) (detailed equations in appendix A). Importantly, the statistical dispersion defined in this statistical framework is not spuriously linked to the number of species in community, as explained in Appendix A.

Posterior distributions of parameters have been sampled by four independent chains using the *No-U-Turn Sampler* implemented in *stan* through the *R* package *brms* (Bürkner, 2017). Careful attention has been paid to built-in diagnostics to avoid divergent iterations and ensure chains convergence. Then, every chain and every posterior distribution have been checked visually, and visual posterior-predictive checks have been performed to ensure that models captured features of the data (Appendix A).

Models have been compared by the mean of weights based on the stacking of predictive distribution. With this method related to Bayesian model averaging, model weights were estimated to maximize leave-one-out predictive density of a complete model containing all sub-models (Yao *et al.*, 2017). The higher the weight of a model was, better were the predictions of future data. It represents one of the less biased and less sensible to overfitting method in Bayesian model selection, and includes uncertainty about every model during weights computation (Yao *et al.*, 2017).

To explore the directionality and intensity of filtering at the community level along the diversity gradient, we used all available data (*n = 2480*). The most complex model (M3d) describing the distribution from which the trait value of the *ith* individual, *y*_*i*_, is drawn, was written as follows:

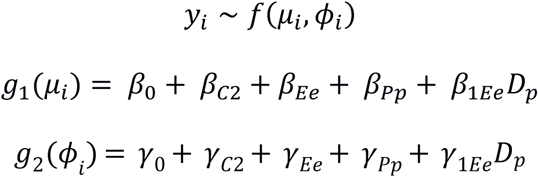

*f()* is a probability distribution parameterized in term of *μ* and ϕ, while *g1()* and *g2()* are link functions. *β*_0_ and *γ*_0_ are intercepts for the first sampling campaign, while *β*_*C2*_ and *γ*_*C2*_ are the deviations for the second sampling campaign for the mean and the dispersion of the distribution, respectively. *β*_*Ee*_ and *β*_*Pp*_ are deviation parameters describing how the mean of each ecosystem *e* and plot *p* differ from the overall mean of each campaign. *γ*_*Ee*_ and *γ*_*Pp*_ are intercepts describing the differences of dispersion. *β*_*Pp*_ and *γ*_*Pp*_ are treated as hierarchical parameters, normally distributed with estimated variances. *β*_*1Ee*_ and *γ*_*1Ee*_ are the ecosystem-specific slopes describing the effect of an increase of one species (D standing for species richness) on the mean and dispersion of community trait distribution, respectively.

To describe the consequences of a variation in species diversity on community trait distribution, we put four models in competition. The reference model, Mcom0, described trait distributions of each plot as a series of intercepts. Mcom1 included a slope per ecosystem describing the link between species diversity and the mean of the community trait distribution, while Mcom2 included a slope linking species diversity to trait dispersion. Mcom3 included both slopes, assuming that biotic filtering exhibited both directionality and varying intensity.

To explore the importance of within species filtering to cope with within ecosystem diversity gradient, we refitted Mcom3 by replacing observed values by mean species value (Mcom3’), estimated from a model containing an intercept for each campaign and a hierarchical intercept per species, distributed normally with estimated standard deviation. By removing intra-specific variation, we aim to create a model with parameters estimated as if the response of community trait distributions were only due to abundance variation. We then evaluate the predictive ability of these model by using observed data. That way, if the predictive ability of this model is lower than models containing ITV, it would mean that abundance variation is not able to capture the process underlying within ecosystems variation in trait distributions.

To analyze consequences of the number of species in community on species trait distribution, we used a subset of data, containing only the individuals of 11 species (*n = 1045*). To be selected, these species had to be dominant when they occurred in poorly diverse plots and growing in communities along the entire diversity gradient of their ecosystem. The most complex model determined both niche mean and dispersion as a function of intercepts and diversity, with equations for *μ* and ϕ:

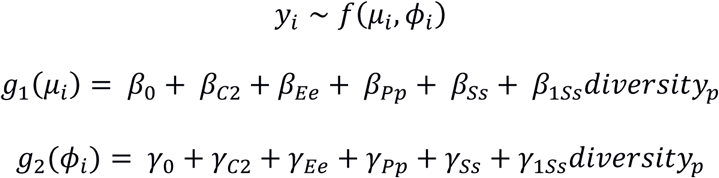

With *β*_*Ss*_ and *γ*_*Ss*_ being species-specific deviation parameters for mean and dispersion, and *β*_*1Ss*_ and *γ*_*1Ss*_ species-specific slopes between species richness and mean and dispersion, respectively. They are all hierarchical parameters distributed multinormally with estimated covariance matrix.

The reference model, Msp0, contained only the series of intercepts, while Msp1 contained a slope per ecosystem linking both species mean and dispersion to species richness. Msp2 included a slope per ecosystem for dispersion, but a slope per species linking diversity to species mean trait value. The more complex model, Msp3, allowed mean and dispersion of each species to move idiosyncratically with the number of species with which they grow.

To explore the strategies of species to cope with variation in biotic conditions, and their potential impact on fitness, we summarized for each species the traits and the density responses. Given the importance of vegetative reproduction in wetlands, we considered density as a good proxy of species demographic performance (7.9% of our individuals were harvested with flower or fruits). To explore responses of species relative density along the diversity gradient, we estimated a hierarchical model including a series of intercepts for campaigns, ecosystems and species identity, and a slope per species linking taxonomic diversity of each plot to the relative density of the species of interest. Species intercepts and slopes were distributed multinormally with estimated covariance matrix. Adapting the framework presented by (Richards *et al.*, 2006), we considered a dominant species with high density in poor communities and low density in rich communities as exhibiting a “master of some” strategy, with plasticity used to take overwhelming advantage when in favorable conditions. A plastic species maintaining its density equal all along the diversity gradient was considered as a “jack-of-all-trades” strategist, potentially using plasticity to cope accurately with multiple conditions.

## Results

### Importance of ITV in community responses

*While filtering among ecosystems sorted species to produce an almost complete turnover, filtering of individuals within species were the principal mechanism along the diversity gradient. Both mean and dispersion of traits distribution varied among ecosystems for every trait but LDMC, with the best models being the one containing intercept for each ecosystem for both aspects of trait distribution (Figs. 1a, 1b, Tab. S1). Within ecosystems, estimating the response of community distributions while using species mean instead of observed values always produced worst predictions than estimating community responses including ITV (Table 1, max weight of Mcom3’ = 0.04). Community level response to the environmental gradient among ecosystems*

**Table 1:** Stacking weights of competing models describing trait variation across biological organization levels. *Community level (Mcom)*: *Mcom0* contained only intercepts for campaign, ecosystems and plots identity (the last two considered as a hierarchical parameter with estimated variance). *Mcom1* and *Mcom2* estimated a slope per ecosystem between species richness and mean and dispersion of trait distribution, respectively. *Mcom3* estimated the ecosystem specific slopes for both mean and dispersion considering estimated species mean during each campaign, only, i.e. without considering species plasticity. *Mcom3’* is based on Mcom3 but considering species phenotypic and ontogenic plasticity plasticity between individuals and campaign. *Species level (Msp): Msp0* contains only categorical effects of campaigns, ecosystem, plot and species identity (the last two considered as a hierarchical parameter with estimated variance). *Msp1* is a model containing categorical effects and a slope per ecosystem linking species diversity to traits mean and diversity of each dominant species within this ecosystem. In *Msp2* trait mean is modeled with a slope per species and dispersion per ecosystem, while *Msp3* contained a slope per species for both mean and dispersion. Parameters *β* and *ɣ* represent slopes between species richness and mean and dispersion of trait distribution, respectively. Indices *E* and *S* mean that the slopes were estimated for each ecosystems or for each species, respectively. LA = Leaf Area (cm^2^), SLA = Specific Leaf Area (cm^2^ g^−1^), Chl. = superficial chlorophyll content, Angle = Leaf angle (°), LDMC = Leaf Dry Matter Content (g_dry_ g_fresh_^−1^), Flav. = flavonoids.

|  | Models | Parameters | Length | LA | SLA | Chl. | Angle | LDMC | Flav. |
| --- | --- | --- | --- | --- | --- | --- | --- | --- | --- |
| Community level | Mcom0 | Int | 0.01 | 0.09 | 0.03 | 0.05 | 0.45 | 0 | 0.03 |
| | Mcom1 | $\beta_{1E}$ | 0 | 0.2 | 0 | 0.84 | 0 | 0 | 0 |
| | Mcom2 | $\gamma_{1E}$ | 0.01 | 0 | 0.13 | 0 | 0.09 | 0.17 | 0.07 |
| | Mcom3' | $\beta_{1E}, \gamma_{1E}$,<br>no ITV | 0 | 0 | 0 | 0 | 0 | 0.04 | 0.04 |
| | Mcom3 | $\beta_{1E}, \gamma_{1E}$ | 0.98 | 0.71 | 0.83 | 0.11 | 0.47 | 0.79 | 0.86 |
| Species level | Msp0 | Int | 0 | 0.23 | 0 | 0.31 | 0.4 | 0 | 0.55 |
| | Msp1 | $\beta_{1E}, \gamma_{1E}$ | 0.32 | 0 | 0.38 | 0 | 0.24 | 0.8 | 0 |
| | Msp2 | $\beta_{1S}, \gamma_{1E}$ | 0 | 0 | 0 | 0.69 | 0 | 0 | 0 |
| | Msp3 | $\beta_{1S}, \gamma_{1S}$ | 0.68 | 0.77 | 0.62 | 0 | 0.36 | 0.2 | 0.45 |

**Figure 1:**
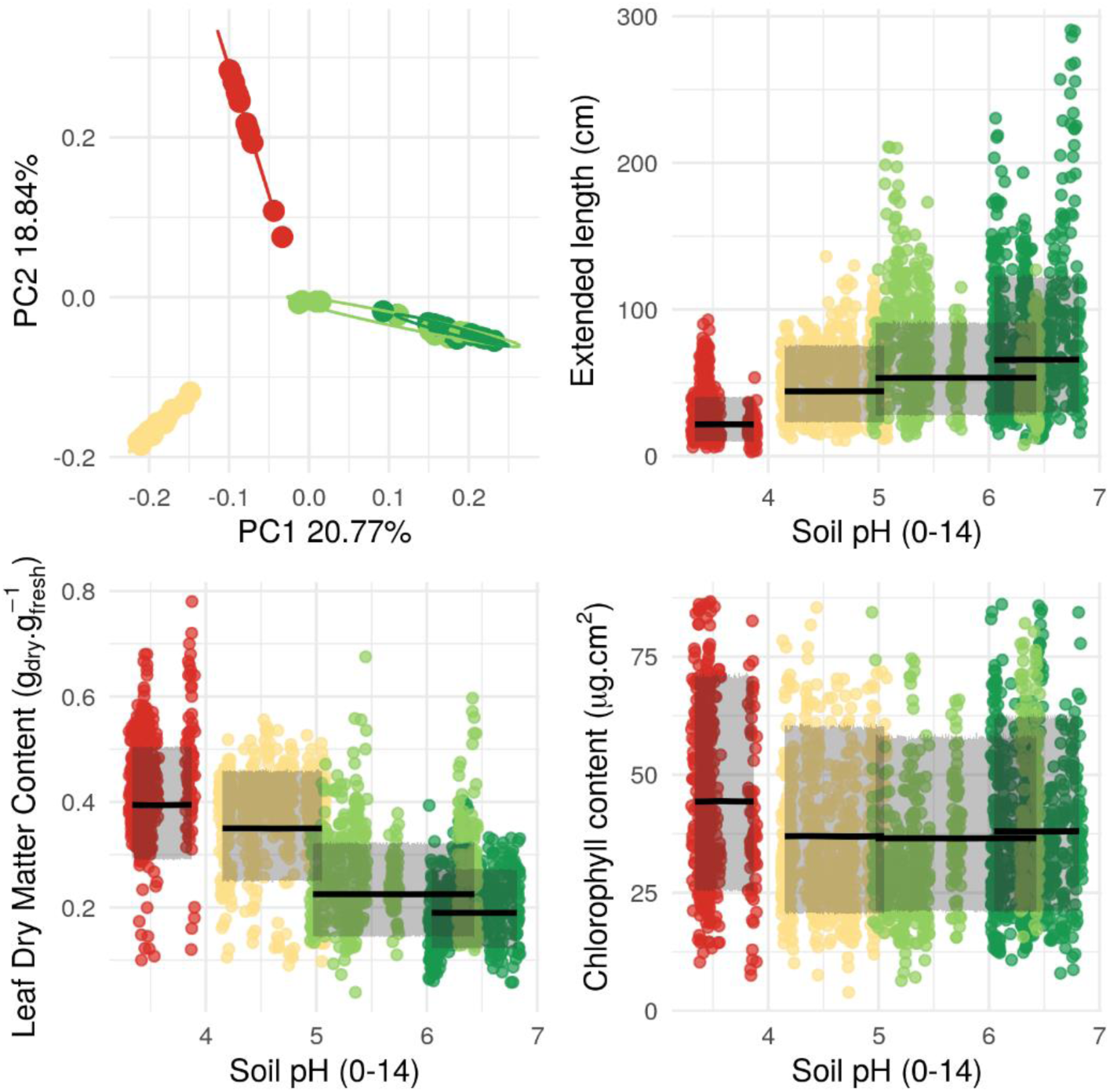
Differences among ecosystems in term of species and functional traits. Panel a represents a principal component analysis of Hellinger transformed species density. Lines represents 95% ellipses capturing 95% of the points. Panel b to d represents observed trait values with predictions of the model containing an intercept per ecosystem for both mean and dispersion. Lines represent mean predicted value. Shaded area represent 95% predictive interval of community trait distribution. Panel b and c represent mean Leaf Dry Matter Content and both Leaf Area mean and dispersion varying monotonically among ecosystems. Panel d representsan absence of among ecosystem monotonic variation for chlorophyll content per unit area of leaves.. *Red* = bog, *yellow* = fen, *light green* = wet meadow, *dark green* = fluvial marsh.

While mean and dispersion of traits related to space filling and light acquisition (EL, LA and SLA) increased monotonically along the pH gradient, the mean of traits related to nutrient conservation (LDMC and flavonoids) decreased in ecosystems with higher pH (Fig. 1, Fig. S4). Dispersion of flavonoid values increased along the pH gradient, while model including a varying dispersion of LDMC distributions among ecosystems was not better than the one with varying means only. Leaf angle and chlorophyll content both varied in mean and dispersion between ecosystems, but without ordered pattern along the pH gradient (Fig. S4).

### Community level response to the plant diversity gradient within ecosystems

We detected community level responses to diversity for both mean and dispersion for functional traits in every aspect of plant strategies (Fig. 2). Light interception traits (EL and LA), SLA, leaf angle and nutrient conservation traits (LDMC and flavonoids content) were all best predicted by modelling mean and dispersion (Tab. 1). Chlorophyll was best predicted by modelling only the mean parameter (Tab. 1).

**Figure 2:**
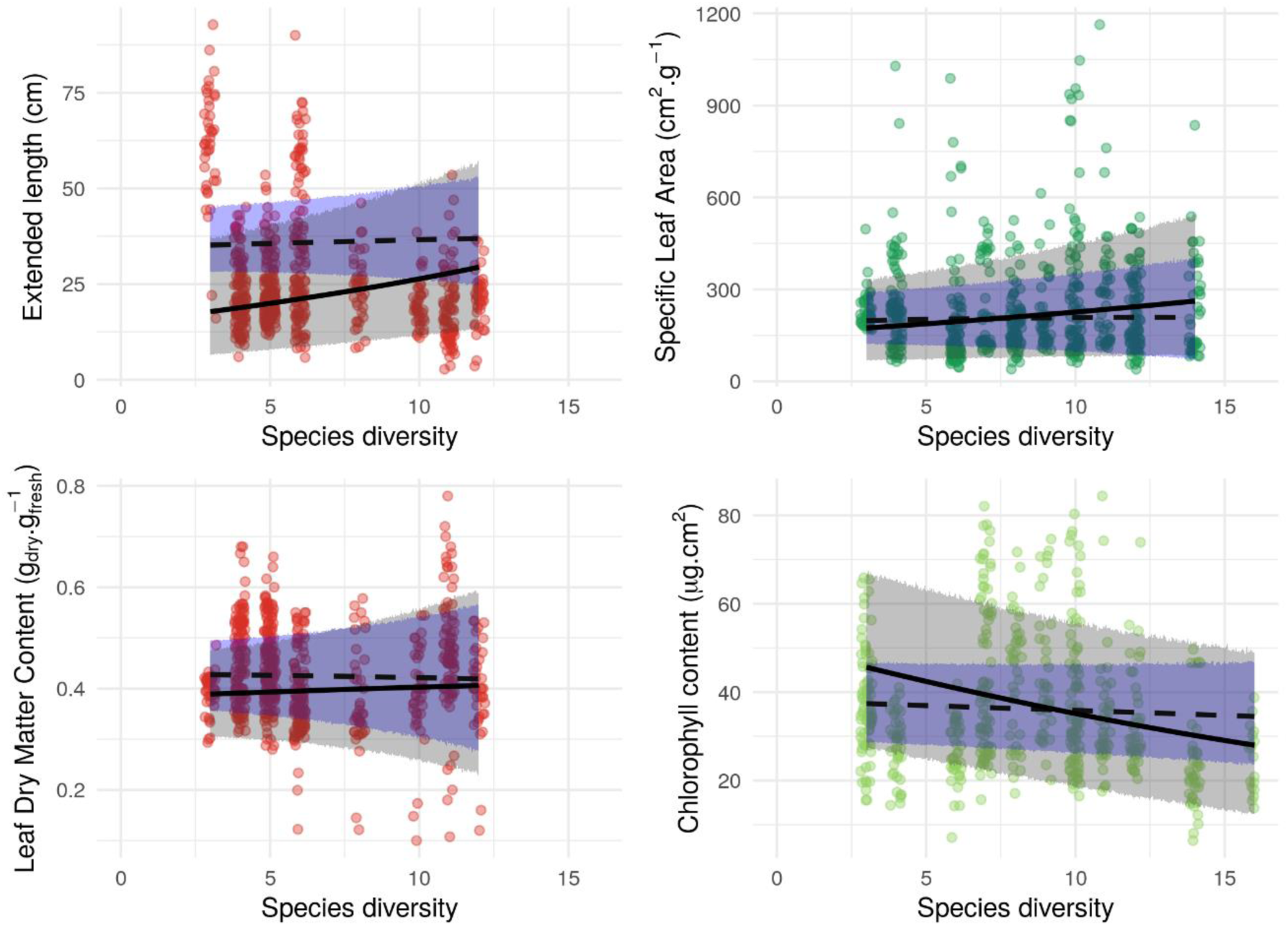
Example of response of community trait distribution to within-ecosystem species diversity gradient. Plain lines represents mean predicted values based on observed data (with ITV), while dashed lines represents mean predicted values of models fitted using species mean values (without ITV). Gray and blue shaded area represents 90% predictive interval of community trait distribution with and without ITV, respectively. *Red* = bog, *yellow* = fen, *light green* = wet meadow, *dark green* = fluvial marsh.

Community functional distributions followed both general and system specific responses. Mean of traits related to space and fine scale light utilization responded in every ecosystem but the wet meadow, suggesting that the number of species in communities imply a directional filtering toward particular values. Both mean and diversity of SLA increased with species richness in every ecosystem, suggesting a great importance of fine scale light conditions and growth trade-offs (Tab. 2, Figs. 2b, S6). In the harshest ecosystems, the bog and the fen, plants were taller in more diverse communities, but with lower LA, while having more dispersed trait values on both traits (Figs. 2a, S4, S5). Interestingly, their leaves presented less chlorophyll per surface, while being also more variable (Fig. S7). LDMC and flavonoids distributions were more variable in more diverse communities only in these ecosystems, suggesting that assemblage of more diverse communities filtered for different strategies of nutrient conservation (Figs. 2c, S9). In the wet meadow, only SLA responded to the diversity gradient, suggesting a lower importance of space, light and nutrient acquisition (Fig. S6). In the fluvial marsh, plants were both smaller and with smaller leaves in more diverse communities, but also more variable (Figs. 2b, S4, S5). Richer communities presented also more variable chlorophyll content and leaf angle distribution, suggesting canopy partitioning and adjustment to fine scale light conditions.

### Species level response to the plant diversity gradient within ecosystems

We detected in every ecosystem two contrasted species responses to the diversity gradient. The first, “Jack-of-all-trade” strategy consisted in species with fixed or slightly increasing median relative density all along the diversity gradient, but with deformed trait distributions (Fig. 3, left part of the panel). This deformation occurred both on mean and dispersion of light and space acquisition related traits, with the best out-of-sample predictions of EL, LA and SLA provided by the model with species-specific slopes within each ecosystem (Tab. 1). For example, *Typha latifolia* exhibited lower and less variable LA in richer fluvial marsh communities, with a median leaf measuring 73.9 (±45.4) cm^2^ when growing with 2 species and 56.8 (±30.1) cm^2^ when growing with 14 species (Fig. 3). However, in the wet meadow, *Lythrum salicaria* trait mean and dispersion and density remained unchanged along the diversity gradient. Conversely, “Master of some” strategies were detected when dominant species were highly dense in taxonomically poor communities, but were less dens in richer communities. Species showing density response may or may not have been plastic responses depending on the trait but tended to present more dispersed trait distribution. For example, in the fen, there was in average 640 (±70) *Carex oligosperma* individuals per m^2^ in 5 species communities, and 66 (±4.4) individuals in 14 species communities. Interestingly, *C. oligosperma* LA, SLA, LDMC and leaf angles were more variables in richer communities, while chlorophyll was less variable. It is worth noting LDMC and chlorophyll species dispersion were best predicted by a common slope shared by species within each ecosystem (tab. 1).

**Fig. 3:**
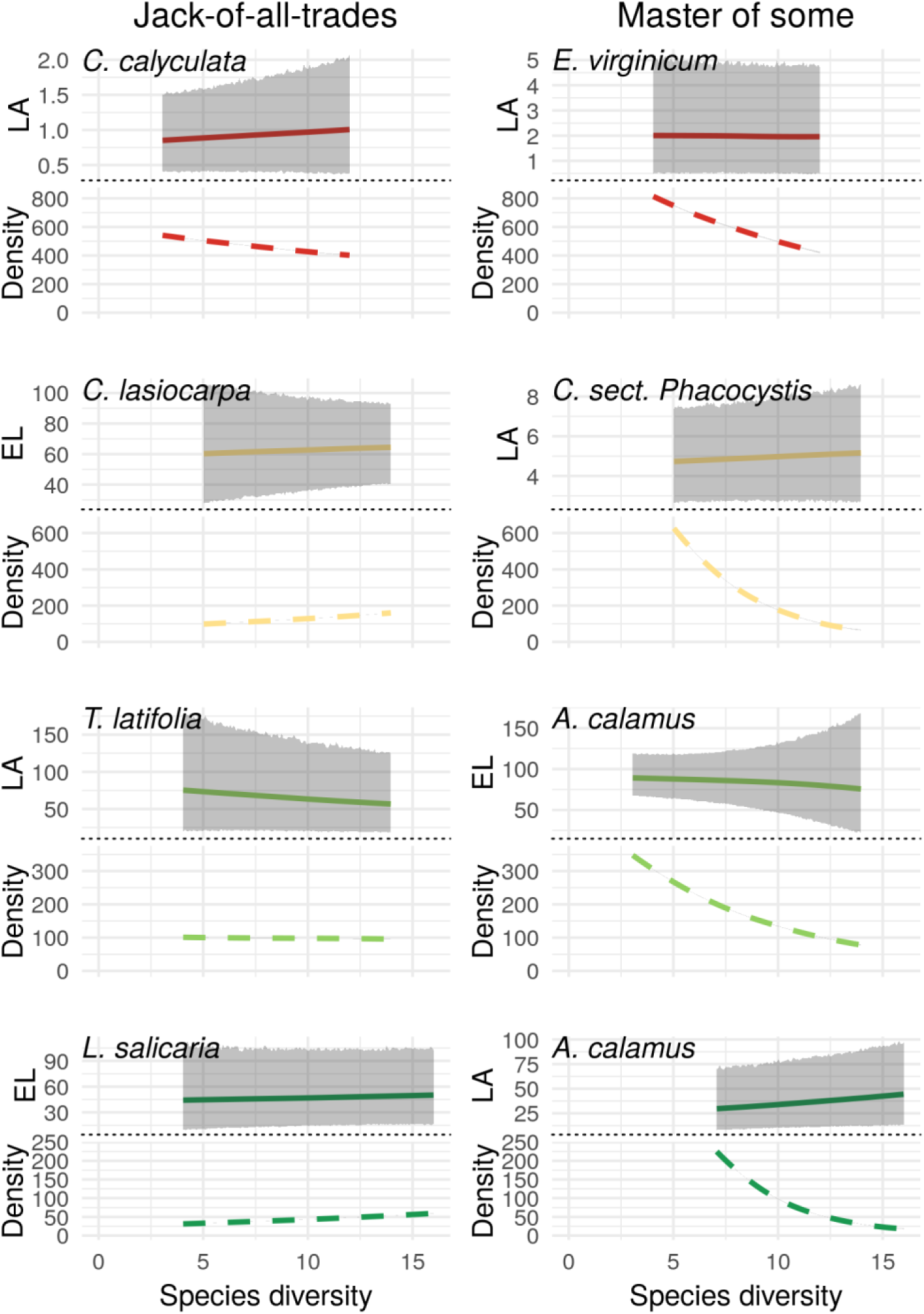
Models predictions of species morphological trait distribution and fitness trait (relative density) in function of the number of species in community, representing the presence of two contrasted response within each ecosystem. The “Master of some” response represents species which were substantially more abundant when they were dominant than when they were growing in rich communities, with or without detected trait variation. This species were potentially able to dominate in favorable conditions, but unable to maintain fitness when conditions changes. The “Jack-of-all-trades” response represents species with stable relative density, but with substantial trait variation, represented by displacement of the mean and/or increase/decrease of diversity of trait values. Potentially, these species were able to cope with changing conditions by deforming their trait distribution. Full line represents the species median trait value, while the grey area represents the 90% predictive interval of trait distribution. Dashed line represents the median predicted relative density of each species. LA = Leaf area (cm^2^), EL = Extended length (cm), Density = Relative density (ind.m^−2^). Species are *Chamaedaphne calyculata, Eriophorum virginicum, Carex lasiocarpa, Carex sec. phacocystis, Typha latifolia, Acorus calamus et Lythrum salicaria*.

## Discussion

We detected change to the distribution of functional traits, in both direction and intensity, along a gradient of plant diversity replicated in four highly contrasted ecosystems filtering for different trait distributions. Such biotic filtering structured trait variation at the community and species levels on respective dimensions of plant strategy. Our study reveals that multiple mechanisms detectable at different levels of biological organization are concomitantly at play to shape the distribution of traits along environmental gradients. This echoes recent studies showing that the drivers of trait variations may not be the same while looking within and between species(e.g. Anderegg et al. 2018), or while sampling at different spatial scales (Messier *et al.* 2016). However, our study is the first one to structure the different levels of trait variation within a common statistical framework and to show *in-natura* that the within-species variation can be strongly structured by biotic interactions.

Our results revealed two sets of traits each responding at a specific level of biological organisation: across ecosystems, every trait was better predicted by including ecosystems identity for mean and dispersion, some of them showing directional and intense filtering in line along the soil fertility gradient. Mean traits related to space occupation (EL, LA) and to nutrient acquisition (LDMC and flavonoids content) increased and decrease from less to more fertile ecosystems, respectively. Selection intensity showed an opposite pattern, trait dispersion of EL, LA and flavonoid increasing. Dispersion of LDMC tended to decrease in more fertile ecosystems (Fig S4), but this was not supported by model selection. These results are coherent with known functional trade-off between nutrient conservation and growth along fertility gradients (Jager *et al.*, 2015), along which plants selected in the more fertile ecosystems are the ones with the lowest investment in leaf longevity but with the greatest ability to compete for space (Grime, 1977; Wright *et al.*, 2004). On the contrary, while an almost complete turnover of species drove differences between ecosystems, SLA and fine-scale light acquisition traits (chlorophyll content, leaf angle) did not show clear patterns across ecosystems. Within ecosystem, filtering along the species diversity gradient selected individuals from a common species pool particular to each ecosystem. Filtering intensity was observed with an increase of the dispersion of trait values for traits related to every functional axis. Within species, filtering intensity was particularly associated with EL, LA and SLA, only. This suggests that biotic interactions have particularly constrained species to deform their niches within their phenotypic plasticity to adapt to changes in space and/or light resources.

Communities’ trait distribution revealed both filtering directionality and intensity along the diversity gradient (Table 2). Both in nutrient-poor and in nutrient-rich ecosystems, less diverse communities exhibited mean leaves with less area per gram than in less diverse ones. This reveals that individuals exhibited more long-life, robust and uniform leaves when neighbors were conspecific. Most of the time, these individuals were graminoid-like species with self-supported great leaves (e.g. *Typha* in nutrient rich ecosystem, *Eriophorum* and *Carex* in nutrient-poor ecosystem). With increasing species diversity, individuals were, on average, longer and exhibited, smaller and lower chlorophyll-investment leaves, and were especially characterized by more heterogeneous leaves when neighbors are heterospecific. Such trait response likely resulted from trade-offs in resource allocation to adapt to light conditions: light capture (high SLA, low chlorophyll) vs light use (opposite syndrome). This is well-known that competition for light structures plant communities in nutrient-rich environments but this has rarely been highlighted in resource-poor environments (but see Wiktor and Diggelen 2004). While responding mainly to the abiotic gradient across ecosystems, LDMC showed also a biotic filtering which was more intense in less diverse communities, particularly in acidic ecosystems. While the statistical dispersion of LDMC values increased with diversity in more acidic ecosystems, the mean remained stable. This suggest that a limit-to-similarity process takes place in these ecosystems relatively to the way nutrients are conserved, coexistence in diverse communities being based upon a long-term partitioning of soil resources (McKane *et al.*, 2002; Gubsch *et al.*, 2011), with a non-detectable selection for a value conferring disproportional fitness advantage.

**Table 2:**
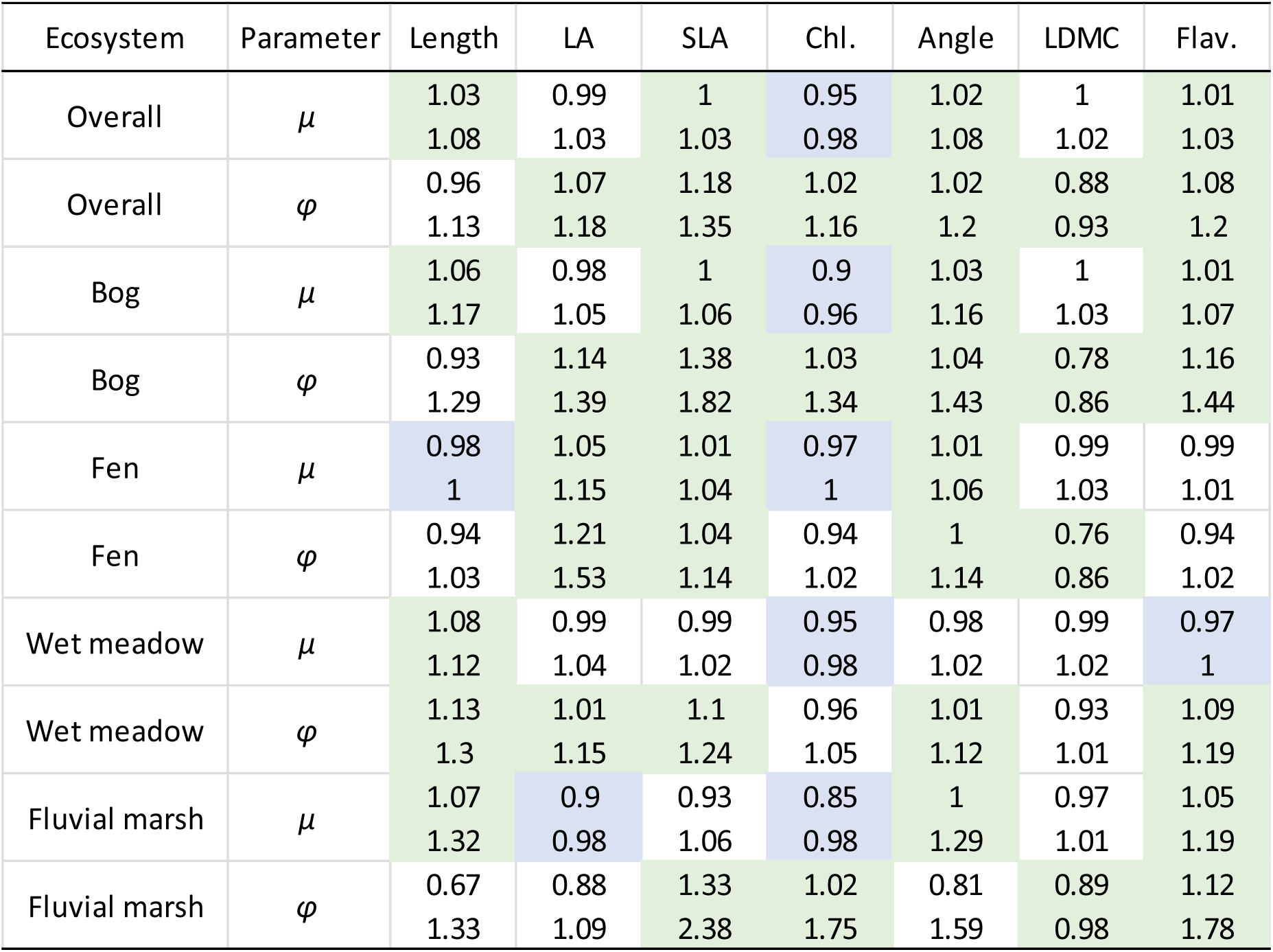
Summary of estimated communities’ trait distribution multiplicative slopes with species diversity. The overall lines represent the global mean response across the data set. We displayed 80% credible intervals of exponentiated slopes. Values before 1 characterizes negative slopes, while values above one characterizes positive ones. *µ* represents the mean of the distribution, while *φ* represents dispersion. Green and blue case represents negative and positive substantial variation in mean trait value (*µ*) or dispersion or precision (*φ*) in response to variation in community species richness. Predictions are available in Fig. S5-S11. LA = Leaf Area (cm^2^), SLA = Specific Leaf Area (cm^2^.g^−1^), Chl. = superficial chlorophyll content, Angle = Leaf angle (°), LDMC = Leaf Dry Matter Content (g_dry_.g_fresh_^−1^), Flav. = flavonoids.

**Table 3:** Summary of estimated species slopes along the species diversity gradient. Parameters were recovered from the best model for each trait, presented in table 1. We removed models predicting LDMC and flavonoids as they had less support for species specific variations and presented no substantial estimates. For sake of clarity, we resumed slopes by symbols representing their sign. We considered the inclusion of zero in the 85% credible interval of parameters. Given the complexity of estimated models, a more conservative interval would exclude pertinent inferences. For the density, symbols + and – represents substantial variation of species mean relative density with species richness within each ecosystem. For each trait they represented substantial variation in mean trait value (mu) or dispersion (phi) in response to variation in community species richness. Model predictions are available in Fig. S12-18. LA = Leaf Area (cm^2^), SLA = Specific Leaf Area (cm^2^.g^−1^), Chl. = superficial chlorophyll content, Angle = Leaf angle (°).

| Ecosystem | Species | Density | Parameter | Length | LA | SLA | Chl. | Angle |
| --- | --- | --- | --- | --- | --- | --- | --- | --- |
| Bog | *Chamaedaphne calyculata* | 0 | $\mu$ | 0 | + | 0 | 0 | 0 |
| | | | $\phi$ | + | + | 0 | 0 | 0 |
| | *Eriophorum virginicum* | - | $\mu$ | 0 | 0 | 0 | 0 | 0 |
| | | | $\phi$ | 0 | 0 | 0 | 0 | 0 |
| Fen | *Carex lasiocarpa* | + | $\mu$ | 0 | 0 | 0 | 0 | 0 |
| | | | $\phi$ | - | 0 | - | 0 | 0 |
| | *Carex oligosperma* | - | $\mu$ | 0 | 0 | 0 | 0 | 0 |
| | | | $\phi$ | 0 | + | + | - | + |
| | *Carex sect. Phacocystis* | - | $\mu$ | 0 | - | 0 | 0 | 0 |
| | | | $\phi$ | 0 | 0 | + | - | + |
| Wet meadow | *Acorus calamus* | - | $\mu$ | 0 | + | 0 | 0 | 0 |
| | | | $\phi$ | + | 0 | 0 | 0 | 0 |
| | *Lythrum salicaria* | 0 | $\mu$ | 0 | 0 | 0 | 0 | 0 |
| | | | $\phi$ | 0 | 0 | 0 | 0 | 0 |
| Fluvial marsh | *Acorus calamus* | - | $\mu$ | 0 | 0 | 0 | 0 | 0 |
| | | | $\phi$ | + | 0 | 0 | 0 | + |
| | *Comarum palustre* | 0 | $\mu$ | 0 | + | 0 | 0 | 0 |
| | | | $\phi$ | 0 | 0 | 0 | 0 | 0 |
| | *Lythrum salicaria* | 0 | $\mu$ | 0 | - | 0 | 0 | 0 |
| | | | $\phi$ | + | + | 0 | 0 | 0 |
| | *Typha latifolia* | 0 | $\mu$ | 0 | - | 0 | 0 | 0 |
| | | | $\phi$ | 0 | - | 0 | 0 | 0 |

At species level, functional responses to diversity went through the adjustment of both the mean and the dispersion of their trait values. Importantly, we show that idiosyncratic behaviors characterised the species level compared with the community level. These strong responses allow stating that intra-specific variation was a major specific characteristic to deal with biotic constraints. We showed that more important than the extent of this variation in a community, the ability of species to exploit different regions of the functional space is fundamental to allow coexistence. Species particularly adjusted their space and light related trait distributions, suggesting that their position in the canopy or their light interception are strongly related to the number of species with which they grow. In our study, we showed that dominant species might simultaneously present two strategies when growing with a different number of species. The first one is exemplified by *Acorus calamus* individuals which had far more similar LA and EL when dominant than when coexisting with 16 species, and this in two different ecosystems. At the contrary, *Typha latifolia*, which can also form almost monospecific stands and coexists with *A. calamus* in the more diverse communities, is both smaller and less diverse when growing with a higher number of species.

To understand the co-occurrence of both increasing and decreasing dispersion of dominant species trait distribution along the diversity gradient, we turn our attention on coexistence theories considering demography of coexisting species. Commonly, an adjustment of species distributions is expected to limit a species similarity with potential competitors (decreasing dispersion and overlap, Grime 2006, Violle et al. 2012), or to equalize fitness differences by presenting an especially adapted phenotype, with each species exploiting equally the same resources (increasing dispersion and overlap, Le Bagousse-Pinguet et al. 2014). On the other side, phenotypic plasticity globally decreases the likelihood of long term coexistence by decreasing species-level differentiation and favoring abundant competitors (Hart *et al.*, 2016). Based on our results, we argue that instead of being a signature of a given environment, these two processes act simultaneously in communities, but with different consequences depending upon the demographic strategy of coexisting species. This is coherent with population level frameworks concerning the role of phenotypic plasticity. Richards et al. (2006) described a framework opposing “Master of some” species vs. “Jack-of-all-trades” ones. The formers are able to maintain demographic rates across various environmental conditions, while the later used their intra-specific variation to take a disproportional advantage when in a favorable environment. Aggregating the signature of these two strategies in community-level indices, for example by using averaged trait overlap, might avoid detecting the underlying mechanisms of community assemblage. The averaging of contrasted patterns might lead to unclear or even flat patterns of trait variations, and would decrease our ability to understand how individuals are selected in communities, especially if one does not take demography into account. Our multi-level approach was able to detect both Master-of-Some and Jack-of-all-trades demographic strategies and that these strategies co-occurred within a given community and in all ecosystems.

Showing the importance of intraspecific variation to cope with biotic environment and its relation to demographic strategies, we highlight the potential evolutionary perspective of individual filtering during community assembly, and the need to bring concepts of community and population ecology closer. Hennion et al. (2016) showed that biotic filtering, represented as species diversity, was able to persistently alter the amine metabolic profile of a grassland species. Waterway et al. (2016) demonstrated that competitive interactions have driven the historical diversification of coexisting sedge species in fens. While it is regularly argued that the filtering of individuals within community is of evolutionary importance (Post & Palkovacs, 2009), the joint study of population and community levels are rarely crossbred in community and functional ecology (Salguero-Gómez *et al.*, 2018). Here, we show that using a hierarchical approach along an abiotic and an independent biotic gradient, we were able to better understand how traits varied across scales. Filtering directionality and intensity occurred at each level of biological organization but on different dimensions of plant functional strategy: nutrient-acquisition / conservation trade-off across ecosystems, light-acquisition / utilization trade-off across communities, space positioning across species. Filtering intensity was a strong structuring factor across all levels and should be better considered separately at each of these levels rather than considered as a ratio (Violle *et al.*, 2012). It is worth noting that statistical dispersion used to detect filtering intensity is preserved from spurious sampling effect related to the number of species in communities. Appendix C demonstrates that the mean deviation around the mean does not increase systematically with the number of species but becomes more accurately estimated. In summary, disentangling biological organisation levels and considering a rich set of traits representing different niche dimensions, allowed revealing the simultaneous selection pressures acting on individuals. By focusing on individuals, we were able to link within ecosystem population trait dynamics to two contrasted demographic strategies, highlighting the particular importance of intra specific trait variation for community assembly, and the potential evolutionary consequences of fine scale biotic gradients.

## Acknowledgements

We thank Caroline Beaulieu, Ariane Bisson, Antoine Filion, Hugo Germain, Benjamin Gosselin, Annie Picard, Mélodie Plourde, Alexandre Proulx, Joannie Vertefeuille for their technical help during site sampling and laboratory analyses. We thank Marco Rodriguez for his help on statistical analyses and Fernando Maestre for his input. This study was supported by the Natural Sciences and Engineering Research Council of Canada (NSERC-Discovery-2016-05716 and NSERC-Discovery-2016).

