## Supplemental Figures for "From plant populations to communities: using hierarchical trait environment relationships to reveal within ecosystem filtering"

### Suplementary Figures

Lucas Deschamps

8 novembre 2018

#### Data

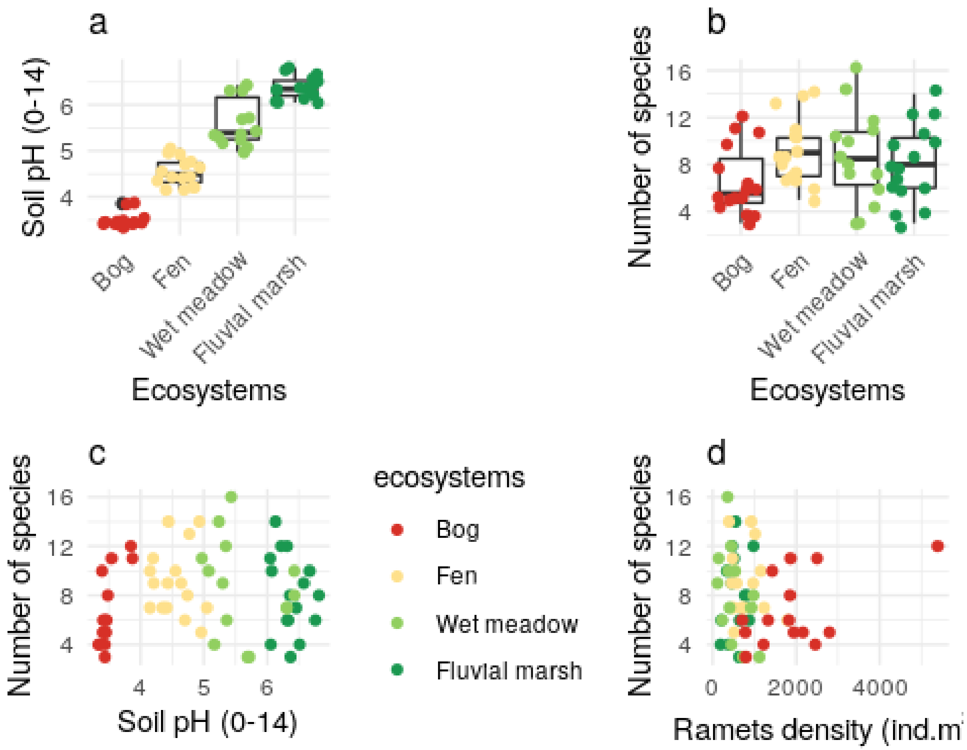


Figure S 1: Abiotic and biotic caracteristics of plots in each ecosystem. Panels a and b present the between ecosystems differences in soil pH and species diversity, respectively. Panels c and d are scatterplots presenting the absence of relationship between species diversity and soil pH and ramets density, respectively.

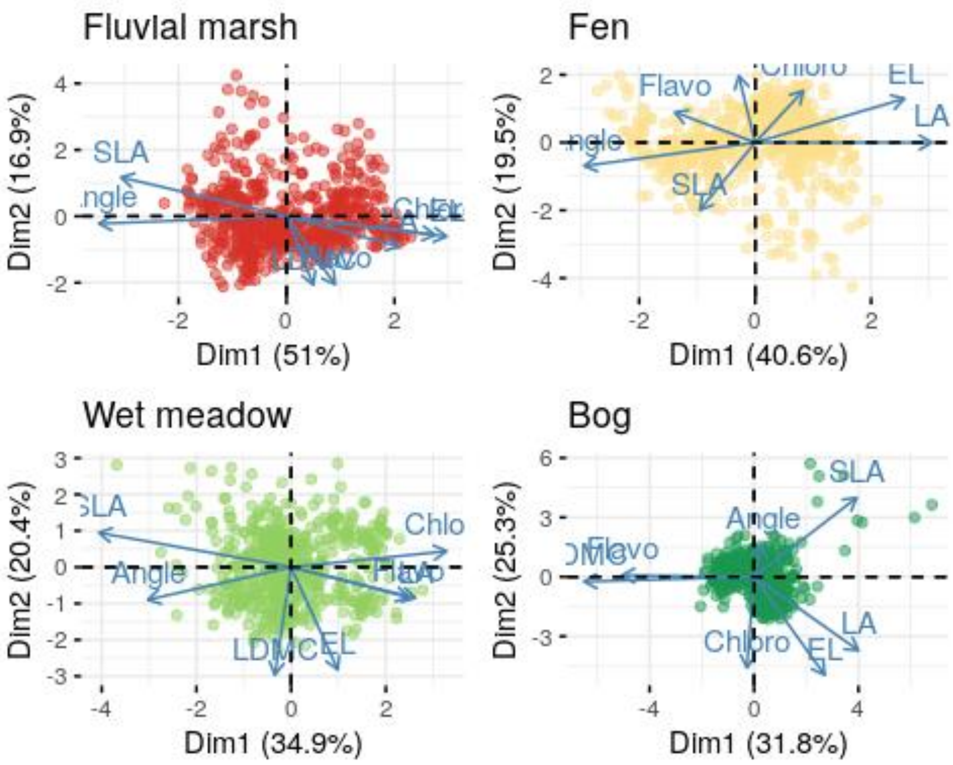


Figure S 2: Varimax rotated principal component analysis of trait values within each ecosystems.

#### Model predictions

##### Ecosystem level predictions

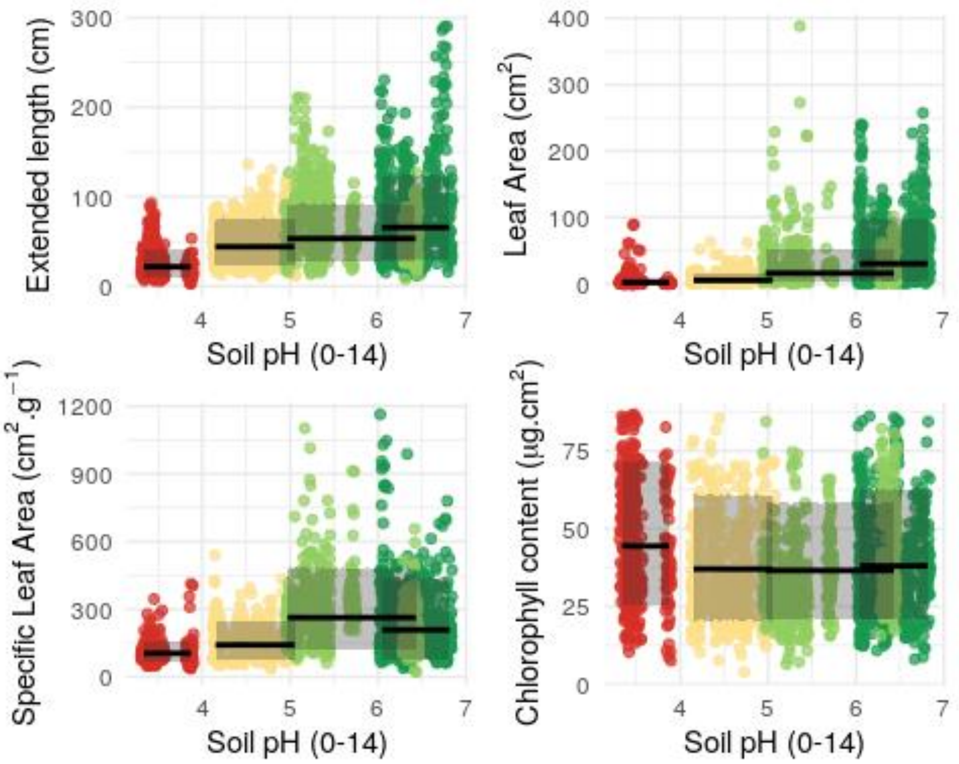


Figure S 3: Between ecosystem differences for the seven studied functional traits. Lines represent the median predicted value, and the shadded areas represent the 80% predictive interval. Red : bog; yellow : fen; light green: wet meadow; dark green: fluvial marsh. Part 1.

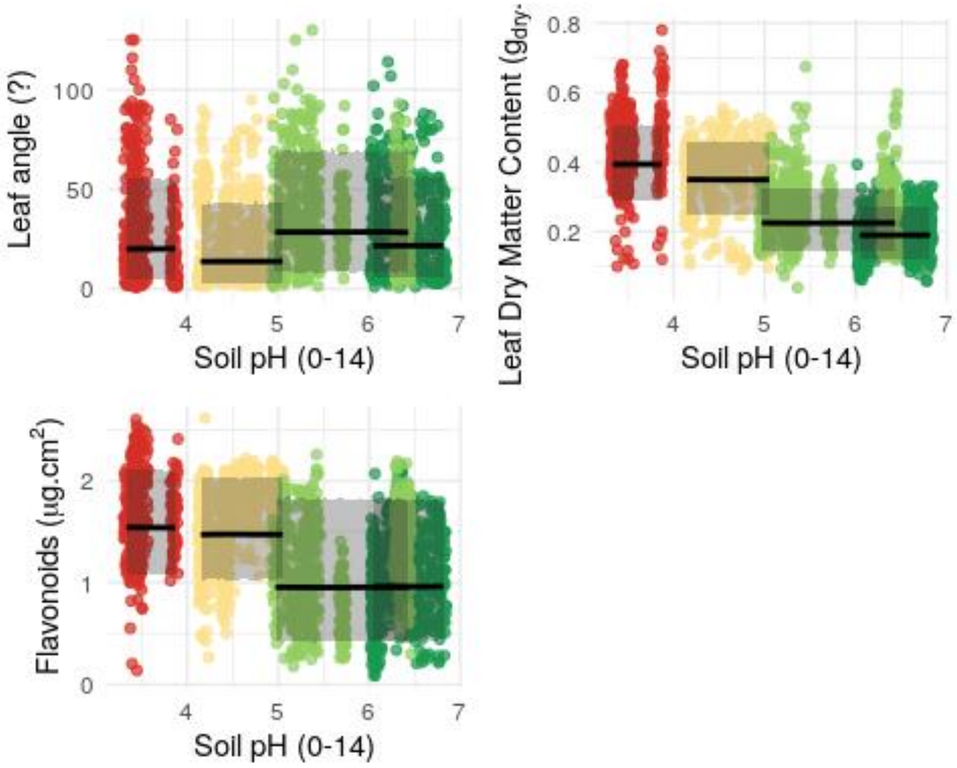


Figure S 3: Between ecosystem differences for the seven studied functional traits. Lines represent the median predicted value, and the shadded areas represent the 80% predictive interval. Red : bog; yellow : fen; light green: wet meadow; dark green: fluvial marsh. Part 1.

#### Community level predictions

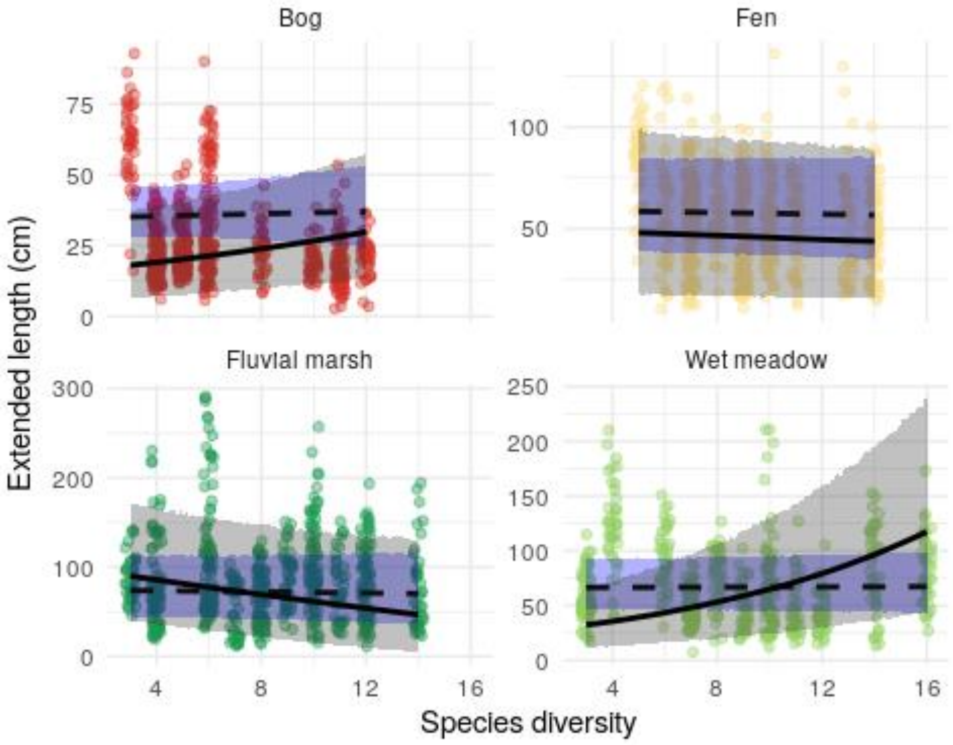


Figure S 4: Relationship between species diversity and extended length community trait distribution in every ecosystem. Lines represent the median predicted value, and the shadded areas represent the 80% predictive interval. Red : bog; yellow : fen; light green: wet meadow; dark green: fluvial marsh.

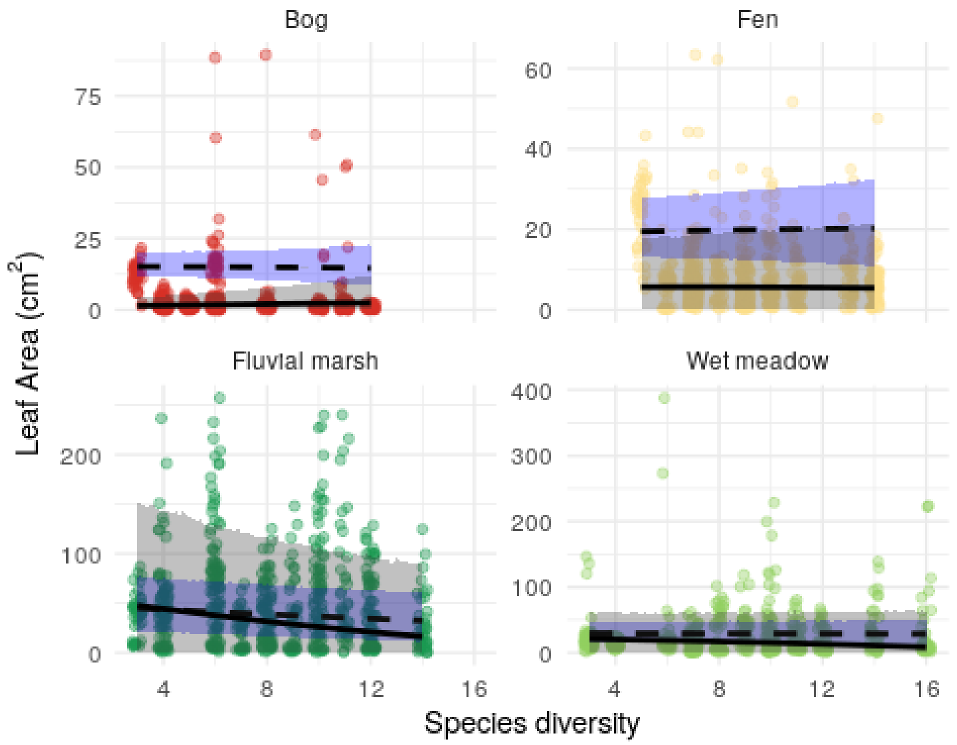


Figure S 5: Relationship between species diversity and leaf area community trait distribution in every ecosystem. Lines represent the median predicted value, and the shadded areas represent the 80% predictive interval. Red : bog; yellow : fen; light green: wet meadow; dark green: fluvial marsh.

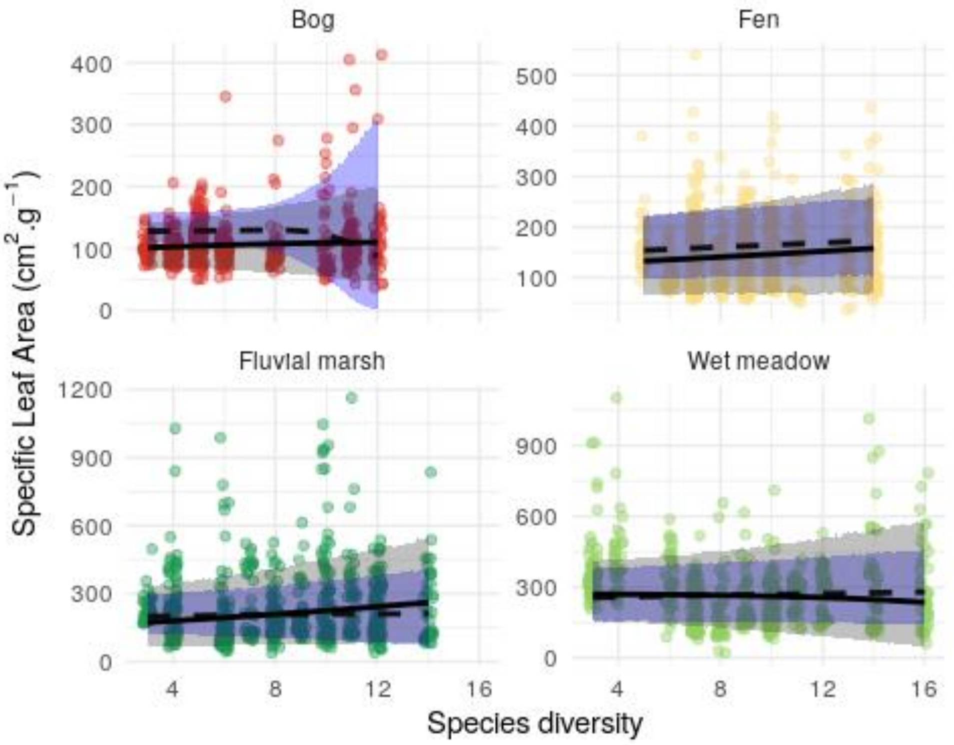


Figure S 6: Relationship between species diversity and specific leaf area community trait distribution in every ecosystem. Lines represent the median predicted value, and the shadded areas represent the 80% predictive interval. Red : bog; yellow : fen; light green: wet meadow; dark green: fluvial marsh.

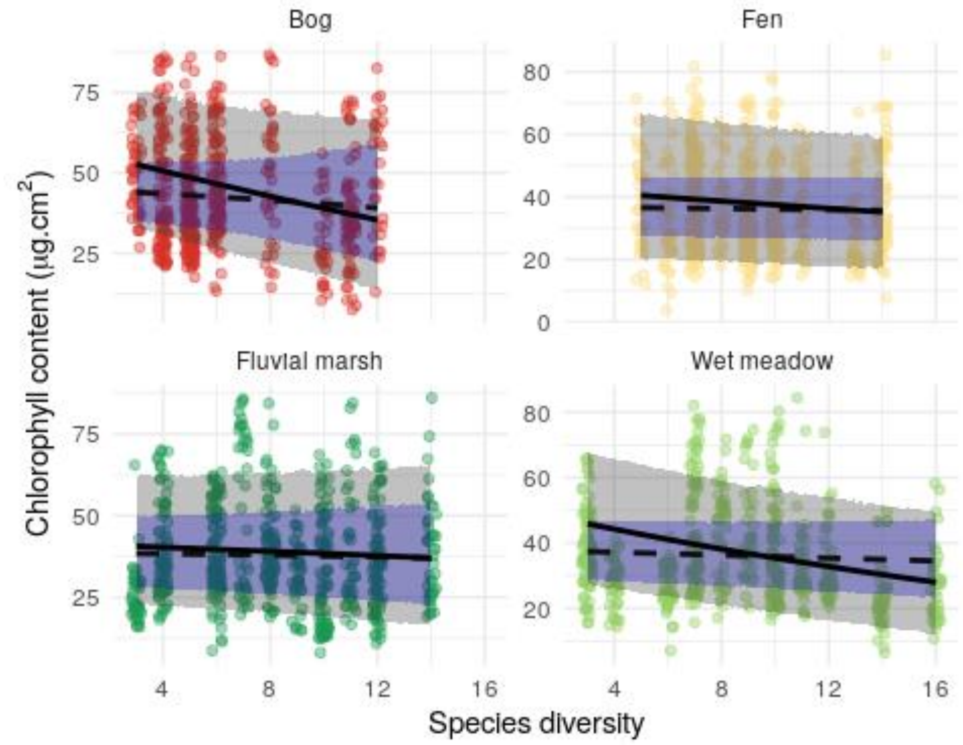


Figure S 7: Relationship between species diversity and superifcial chlorophyll community trait distribution in every ecosystem. Lines represent the median predicted value, and the shadded areas represent the 80% predictive interval. Red : bog; yellow : fen; light green: wet meadow; dark green: fluvial marsh.

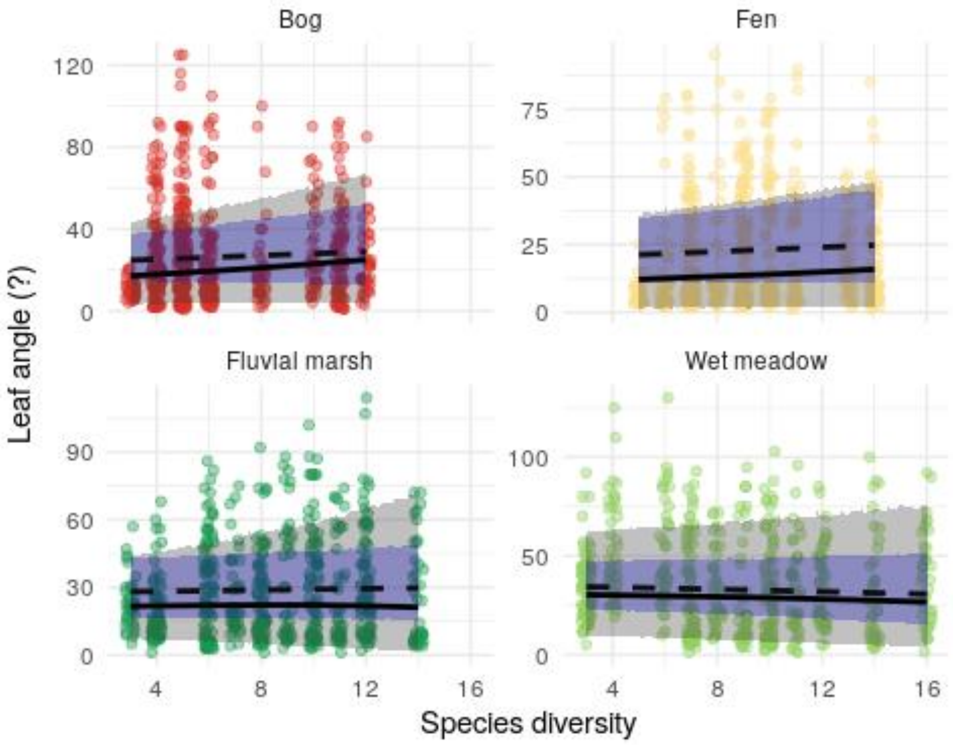


Figure S 8: Relationship between species diversity and leaf angle community trait distribution in every ecosystem. Lines represent the median predicted value, and the shadded areas represent the 80% predictive interval. Red : bog; yellow : fen; light green: wet meadow; dark green: fluvial marsh.

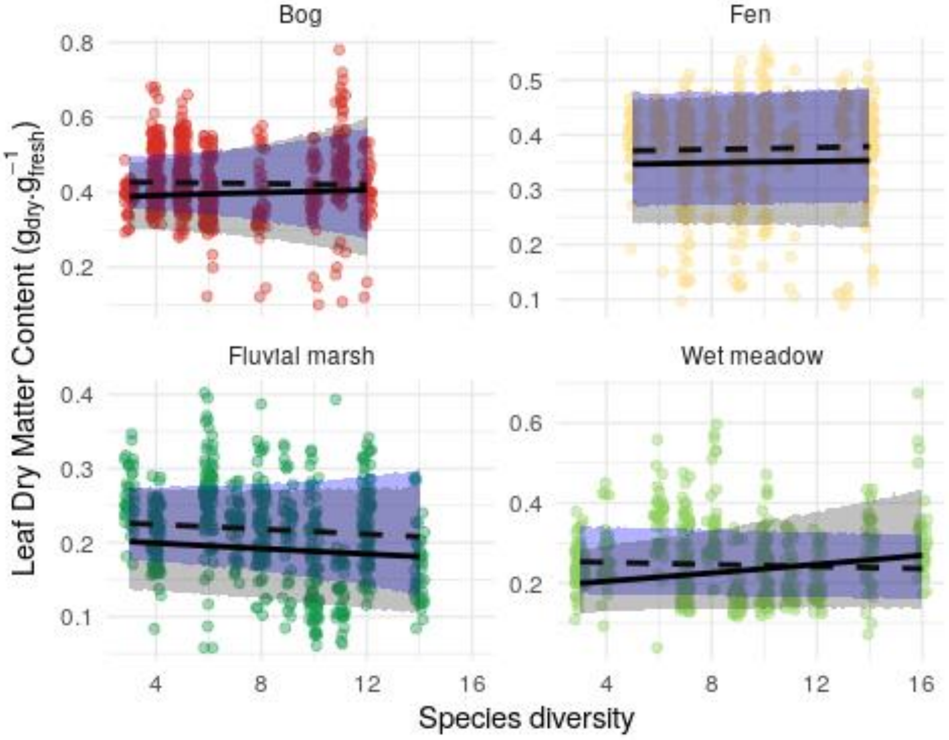


Figure S 9: Relationship between species diversity and leaf dry matter content community trait distribution in every ecosystem. Lines represent the median predicted value, and the shadded areas represent the 80% predictive interval. Red : bog; yellow : fen; light green: wet meadow; dark green: fluvial marsh.

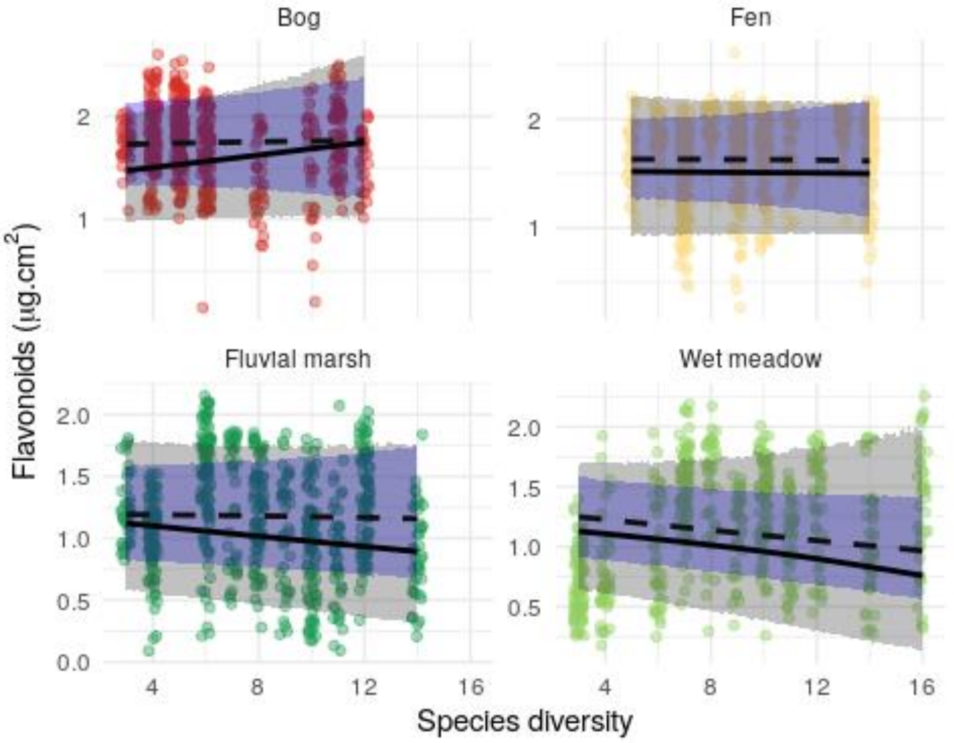


Figure S 10: Relationship between species diversity and flavonoids community trait distribution in every ecosystem. Lines represent the median predicted value, and the shadded areas represent the 80% predictive interval. Red : bog; yellow : fen; light green: wet meadow; dark green: fluvial marsh.

#### Species level predictions

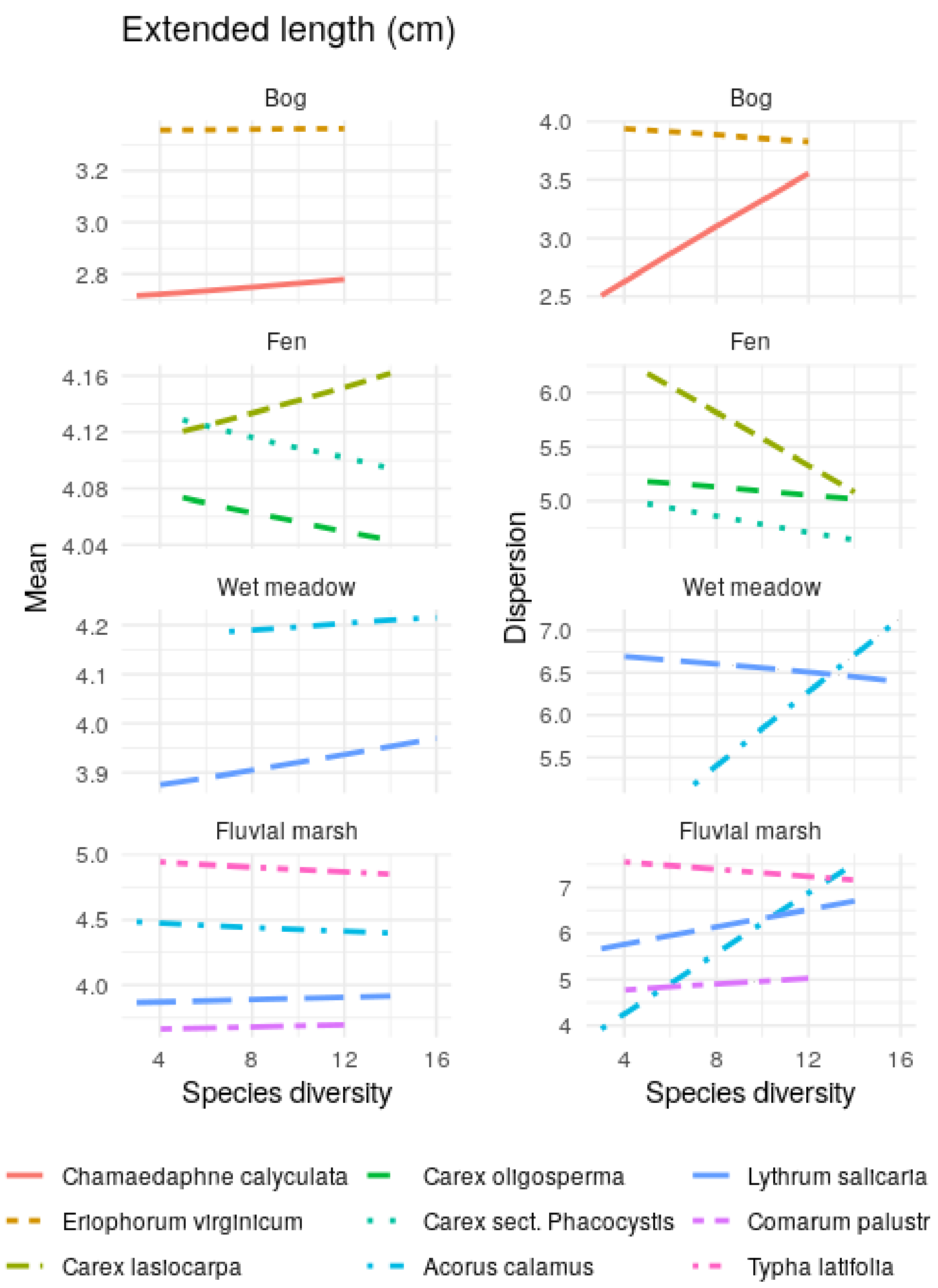


Figure S 11: Relationship between species diversity and species extended length mean value and dispersion in every ecosystem. Lines represent the median predicted value of each parameter. For every trait bu LDMC, phi is a dispersion parameter, while it is a precision parameter for LDMC, modeled with a beta distribution. Red : bog; yellow : fen; light green: wet meadow; dark green: fluvial marsh.

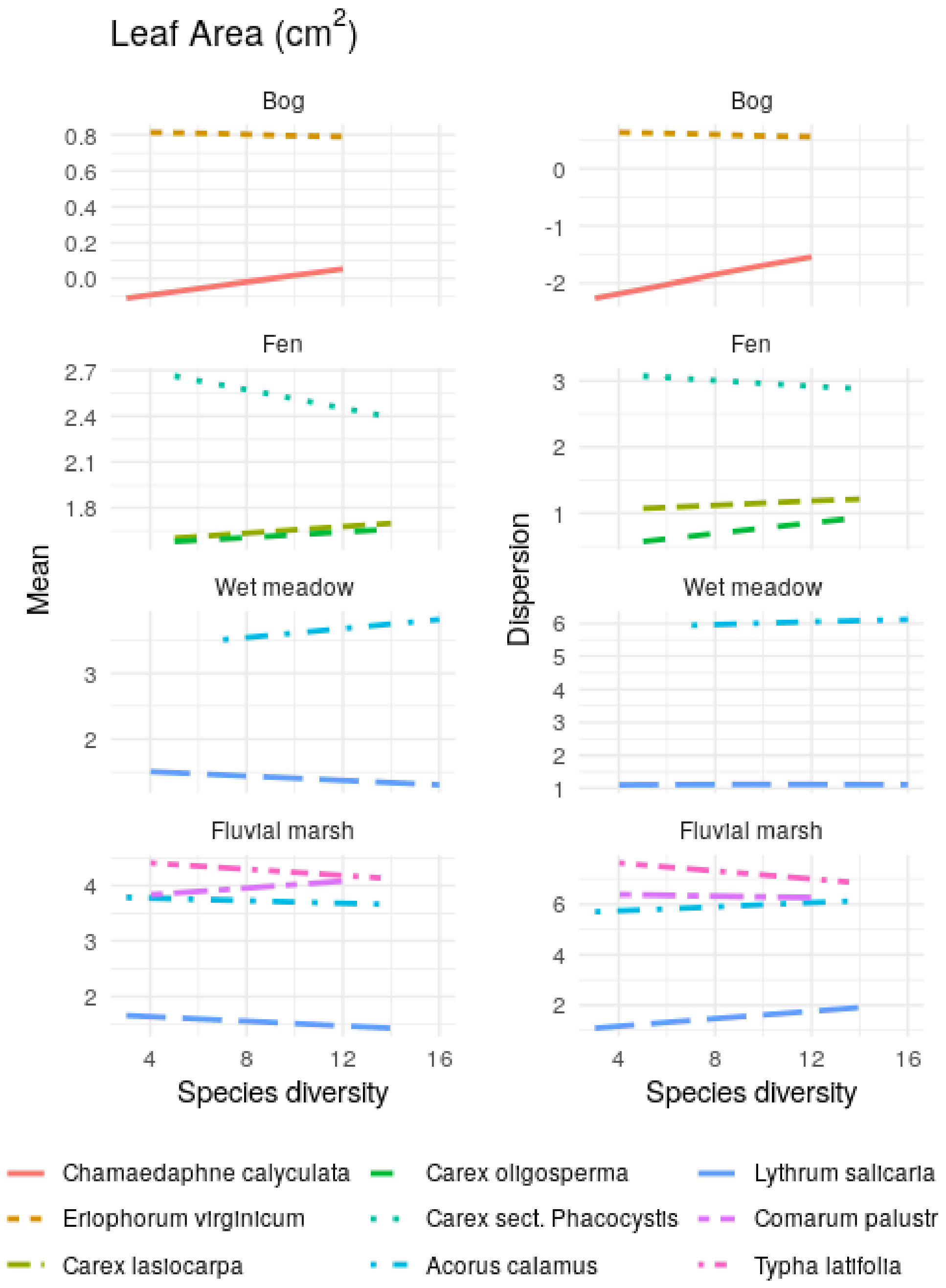


Figure S 12: Relationship between species diversity and species leaf area mean value and dispersion in every ecosystem. Lines represent the median predicted value of each parameter. For every trait bu LDMC, phi is a dispersion parameter, while it is a precision parameter for LDMC, modeled with a beta distribution. Red : bog; yellow : fen; light green: wet meadow; dark green: fluvial marsh.

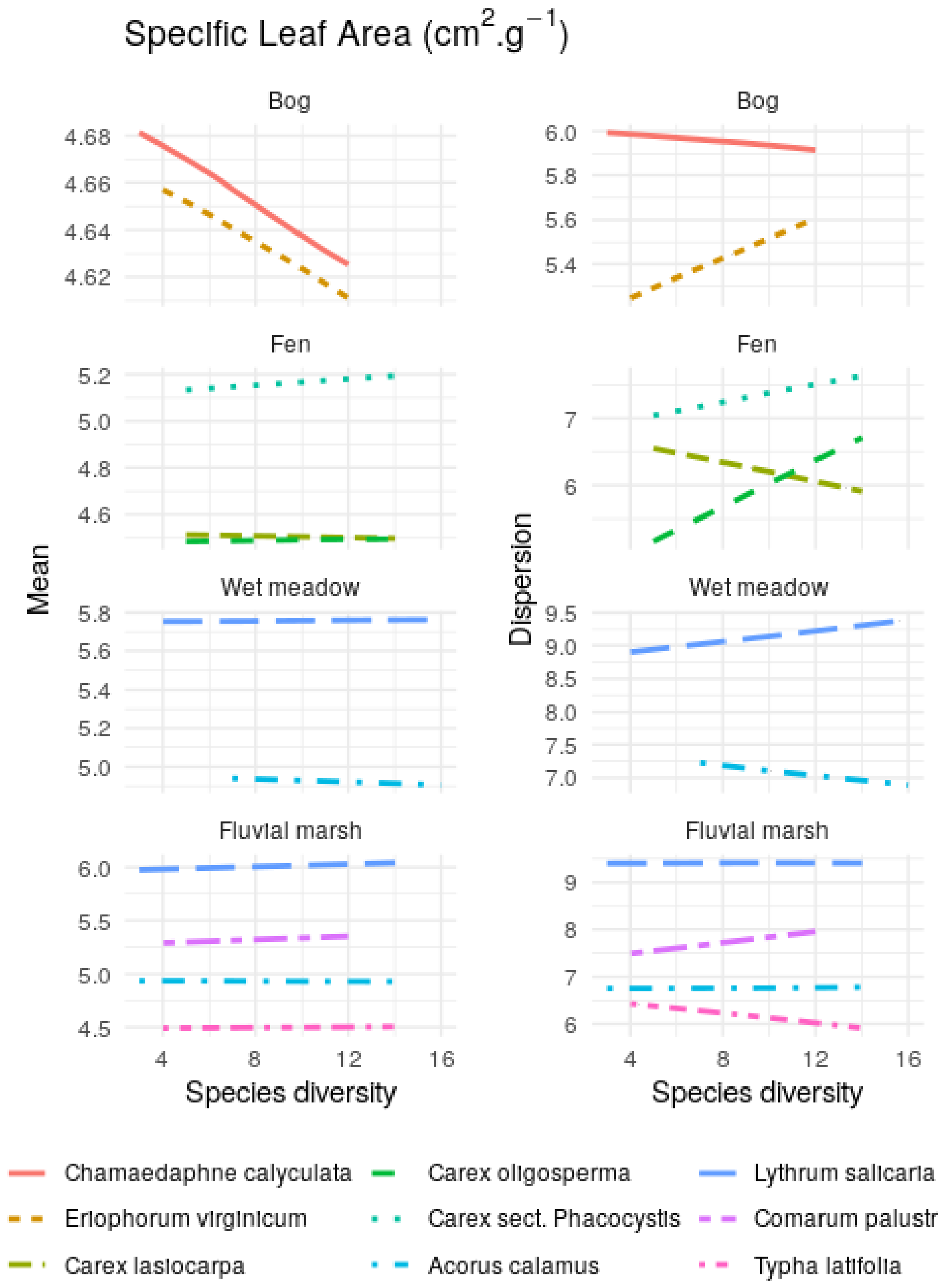


Figure S 13: Relationship between species diversity and species specific leaf area mean value and dispersion in every ecosystem. Lines represent the median predicted value of each parameter. For every trait bu LDMC, phi is a dispersion parameter, while it is a precision parameter for LDMC, modeled with a beta distribution. Red : bog; yellow : fen; light green: wet meadow; dark green: fluvial marsh.

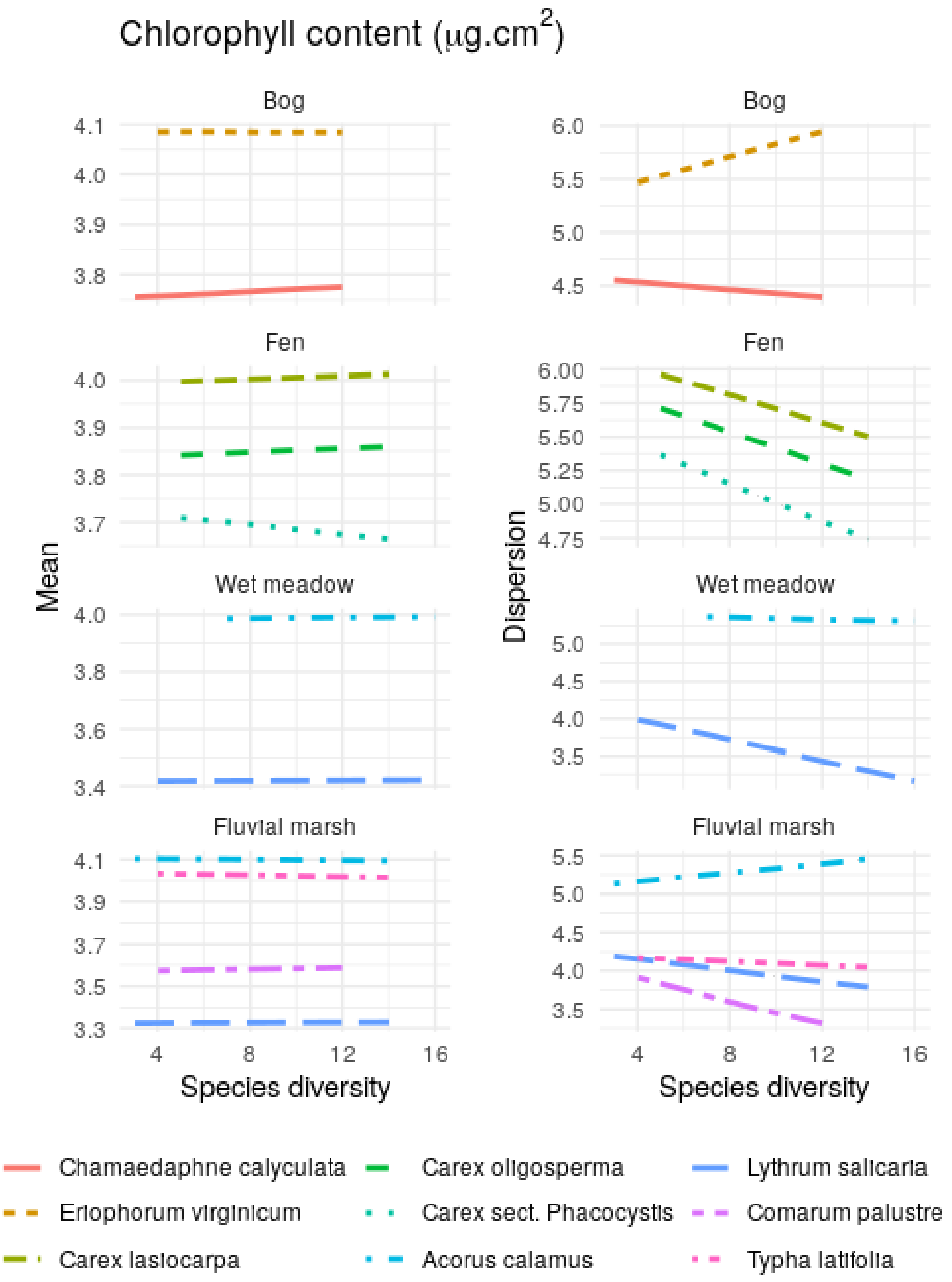


Figure S 14: Relationship between species diversity and species superficial chlorophyll mean value and dispersion in every ecosystem. Lines represent the median predicted value of each parameter. For every trait bu LDMC, phi is a dispersion parameter, while it is a precision parameter for LDMC, modeled with a beta distribution. Red : bog; yellow : fen; light green: wet meadow; dark green: fluvial marsh.

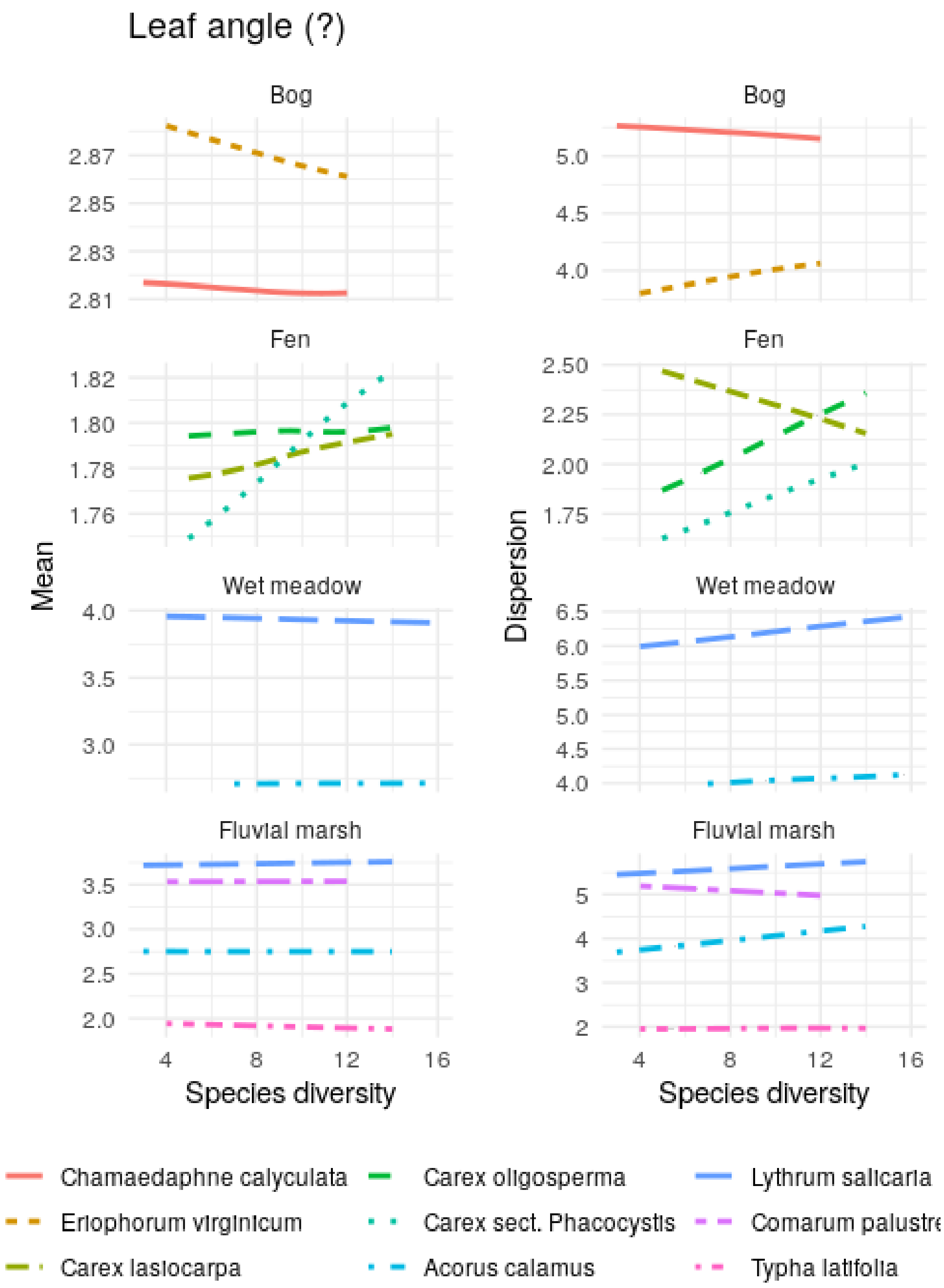


Figure S 15: Relationship between species diversity and species leaf angle mean value and dispersion in every ecosystem. Lines represent the median predicted value of each parameter. For every trait bu LDMC, phi is a dispersion parameter, while it is a precision parameter for LDMC, modeled with a beta distribution. Red : bog; yellow : fen; light green: wet meadow; dark green: fluvial marsh.

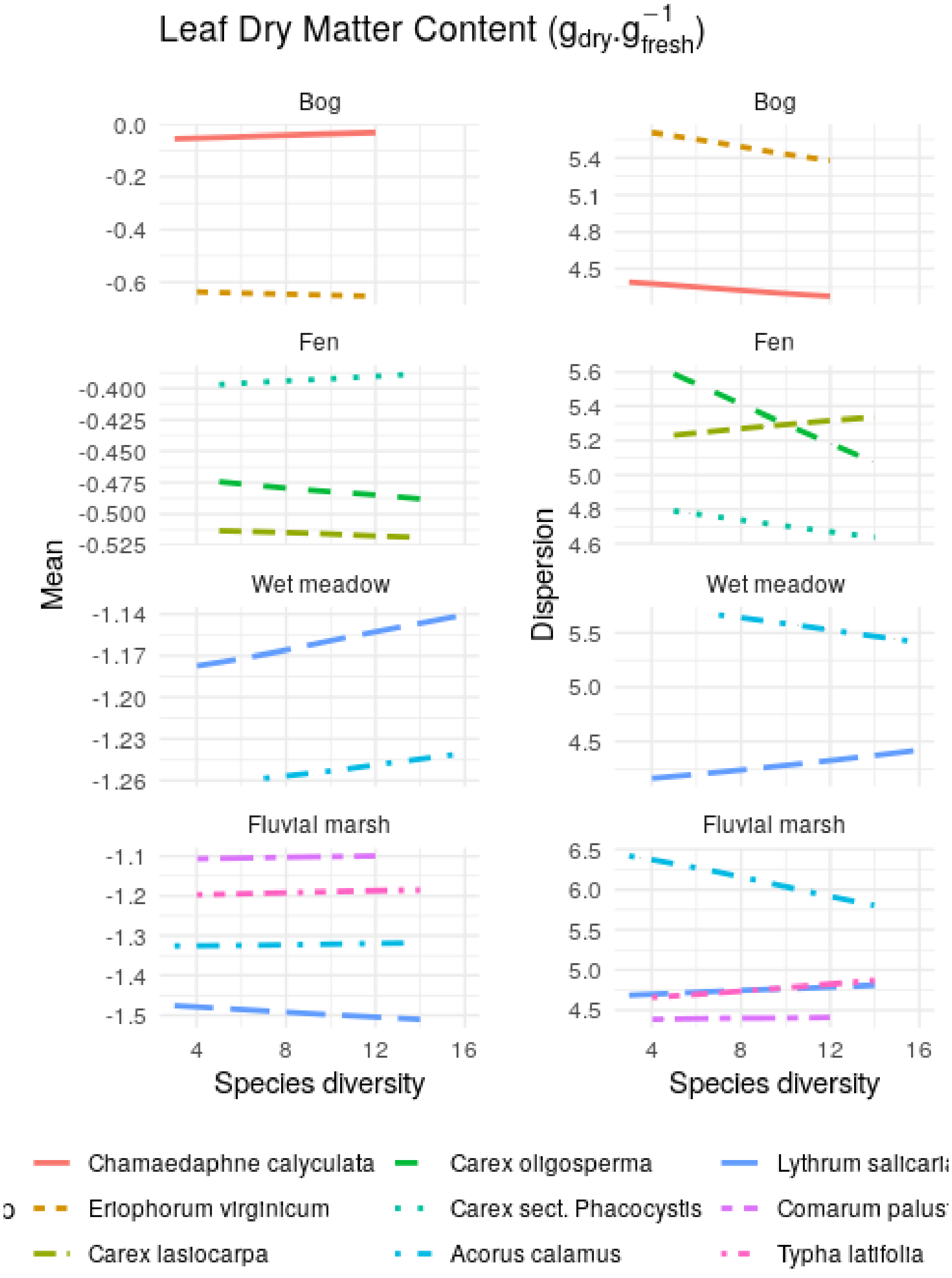


Figure S 16: Relationship between species diversity and species leaf dry matter content mean value and dispersion in every ecosystem. Lines represent the median predicted value of each parameter. For every trait bu LDMC, phi is a dispersion parameter, while it is a precision parameter for LDMC, modeled with a beta distribution. Red : bog; yellow : fen; light green: wet meadow; dark green: fluvial marsh.

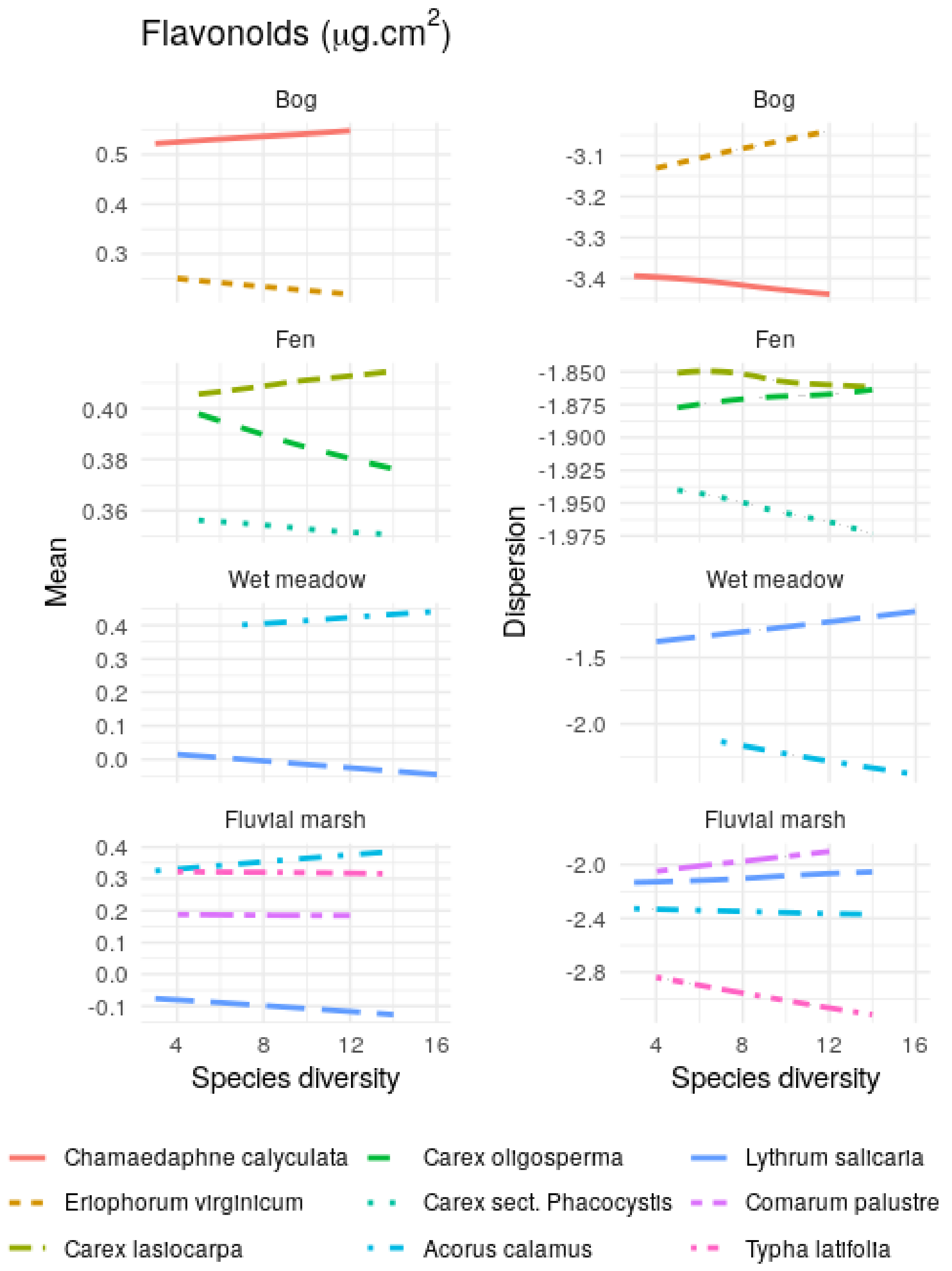


Figure S 17: Relationship between species diversity and species flavonoids mean value and dispersion in every ecosystem. Lines represent the median predicted value of each parameter. For every trait bu LDMC, phi is a dispersion parameter, while it is a precision parameter for LDMC, modeled with a beta distribution. Red : bog; yellow : fen; light green: wet meadow; dark green: fluvial marsh.
